# A PLETHORA transcription factor shapes cucumber shoot architecture

**DOI:** 10.1101/2025.09.18.677013

**Authors:** Merijn Kerstens, Florian Müller, Kelvin Adema, Olga Kulikova, Magdalena Lastdrager, Ben Scheres, Viola Willemsen

## Abstract

- PLETHORA transcription factors (PLTs) are master regulators of plant development. Loss of shoot meristematic PLTs leads to reduced phyllotactic regularity and robustness in Arabidopsis and increased inflorescence branching in tomato. Whether these factors have similar functions in other species is not known.
- To address this knowledge gap, we integrated phylogenetic, transcriptomic, and genome-wide *in vitro* binding strategies with quantitative shoot architecture phenotyping of a panel of cucumber TILLING mutants of each *CsPLT* homolog.
- We determined that *CsPLT3/7* and *CsPLT* are expressed in complementary domains in the cucumber shoot apex. DAP-seq data indicated that both transcription factors recognised an ANT-like consensus motif and bound to SAM organising genes. We identified strong phenotypic defects in multiple *Csplt3/7* mutants. In mature seeds, *Csplt3/7* mutants formed a flat apex instead of a shoot apical meristem, from which secondary meristems emerged during subsequent seedling development. Mature *Csplt3/7* shoots exhibited defects in internode length regularity, stem architecture, flower morphology, and axillary organ initiation. Moreover, phyllotactic patterns were slightly shifted from a spiral towards a distichous orientation.
- We present one of the first pieces of evidence that the PLT3/7 clade fulfils both conserved and diversified roles in shoot development across taxonomic family boundaries.

## Introduction

In plants, the shaping, spacing and orientation of aboveground tissues collectively establish a certain shoot architecture. Shoot growth is a modular process, in which nodes harbouring lateral organs such as leaves, flowers and axillary shoots are separated by organ-free internodes (Wang *et al*., 2018), the total product of which determines optimal cultivation strategies in agriculture. For instance, the economically relevant species cucumber (*Cucumis sativus*) grows long, vining stems that in modern high-wire cultivation methods are guided upwards along strings. Cucumber plants form one leaf per node, while branches, tendrils and flowers emerge from the leaf axils, thus combining reproductive and vegetative growth in a single shoot axis (Liu *et al*., 2021).

Cucumber leaves are formed at the shoot apical meristem (SAM), a dome-shaped structure from which primordia emerge rhythmically. SAM anatomy and primordium initiation have been studied extensively in Arabidopsis (*Arabidopsis thaliana*). In this species, the vegetative SAM and inflorescence meristem (IM) domes can be subdivided into distinct domains. At the dome summit, the central zone (CZ) contains slowly dividing stem cells whose progeny move outward to the peripheral zone (PZ), gradually differentiating to allow production of new organs (Reddy *et al*., 2004). Spatiotemporal integration of auxin signalling is a major factor in specifying the sites at which such organ primordia can be formed, giving rise to highly non-random phyllotactic patterns (Reinhardt *et al*., 2003; Heisler *et al*., 2005; Smith *et al*., 2006; de Reuille *et al*., 2006; Jönsson *et al*., 2006; Galvan-Ampudia *et al*., 2020).

Stem-cell identity of the CZ is maintained by the underlying organizing centre (OC), a group of cells that prevents cell differentiation through a negative feedback loop between the CZ-secreted CLAVATA3 (CLV3) peptide and the OC-produced mobile WUSCHEL (WUS) transcription factor (Laux *et al*., 1996; Mayer *et al*., 1998; Schoof *et al*., 2000; Brand *et al*., 2000; Müller *et al*., 2006). Whereas loss of CLV3 or its receptor CLV1 leads to increased meristem size and fasciation (Clark *et al*., 1993, 1995), *wus* mutants fail to maintain the stem cell pool in the CZ (Laux *et al*., 1996), highlighting the dynamic interplay of the CLV3-WUS module in shaping SAM function. In cucumber, it was shown that the orthologs of WUS and CLV3 antagonistically control carpel numbers, but they are both expressed exclusively in the zone underneath the stem cells (Che *et al*., 2020). This hints that maintenance of cucumber stem cells might be differently regulated compared to Arabidopsis.

PLETHORA (PLT) transcription factors (TFs) of the euAINTEGUMENTA (euANT) family are master regulators of plant development. PLTs are conserved across angiosperms and can be phylogenetically separated into four homology groups, i.e. PLT1/2, PLT3/7, BABY BOOM (BBM)/PLT4 and PLT5, named according to the grouping of the Arabidopsis homologs (Kerstens *et al*., 2020). In this species, PLTs are thought to confer meristematic potential to critical developmental processes, including (lateral) root development, embryogenesis and regeneration, promoting inception and maintenance of growth apices (Aida *et al*., 2004; Galinha *et al*., 2007; Kareem *et al*., 2015; Du & Scheres, 2017; Kerstens *et al*., 2024). In the SAM and IM, PLT3, PLT5 and PLT7 are expressed in partially overlapping domains and promote robustness of the phyllotactic spiral (Prasad *et al*., 2011; Pinon *et al*., 2013; Kerstens *et al*., 2025a). Meristems devoid of all three PLTs remain functional, but exhibit a delayed convergence from a decussate configuration to the golden angle (137.5°) in the rosette. Additionally, *plt3 plt5 plt7* mutants display rare phyllotactic shifts to 180° and 90° angles in the IM, which was linked to reduced auxin and cytokinin signalling, and transport (Prasad *et al*., 2011; Pinon *et al*., 2013; Kerstens *et al*., 2025a). In tomato (*Solanum lycopersicum*), SlPLT3 and SlPLT7 control branching of the inflorescence, suggesting that PLT3/7 TFs are involved in shoot development across eudicots, but regulate different processes (Zebell *et al*., 2025). Outside of Arabidopsis and tomato, however, no role for PLTs in shaping shoot architecture have been described.

In this study, we investigated whether PLTs are also involved in shoot development in cucumber. We uncovered that only two of the four cucumber *PLT*s (*CsPLT*) are expressed in the SAM, of which only *CsPLT3/7* localises to the stem cell niche. Plants containing a loss-of-function *Csplt3/7* TILLING allele lack SAMs in young seedlings, and the combinatorial loss of CsPLT1/2 and CsPLT3/7 proved embryonic lethal. Although spontaneous SAM emergence occurred at the shoot apex in older *Csplt3/7* seedlings, mature shoots exhibited extreme internode lengths, axillary organ outgrowth defects, abnormal flower configurations and disturbed phyllotaxis. We thus establish CsPLT3/7 as a key developmental TF in shaping cucumber shoot architecture and, more generally, demonstrate shared and distinct developmental roles for PLTs across species boundaries.

## Results

### The cucumber genome contains four *PLT* homologs expressed in root and shoot apices

In order to identify shoot developmental roles of PLTs in cucumber, we first generated a phylogenetic tree of the euANT TF family within 13 species of the cucurbit lineage. In contrast to Arabidopsis, which contains six *PLT* homologs divided over four gene clades (Kerstens *et al*., 2020), cucumber and its close relatives harboured only four *PLT* homologs, one in each clade (Fig. 1a,b). Per-clade *PLT* copy number was generally increased to two in the genus *Cucurbita*, which aligns with the predicted whole-genome duplication event in the tribe Cucurbitae (Sun *et al*., 2017; Montero-Pau *et al*., 2018; Barrera-Redondo *et al*., 2019).

**Figure 1.**
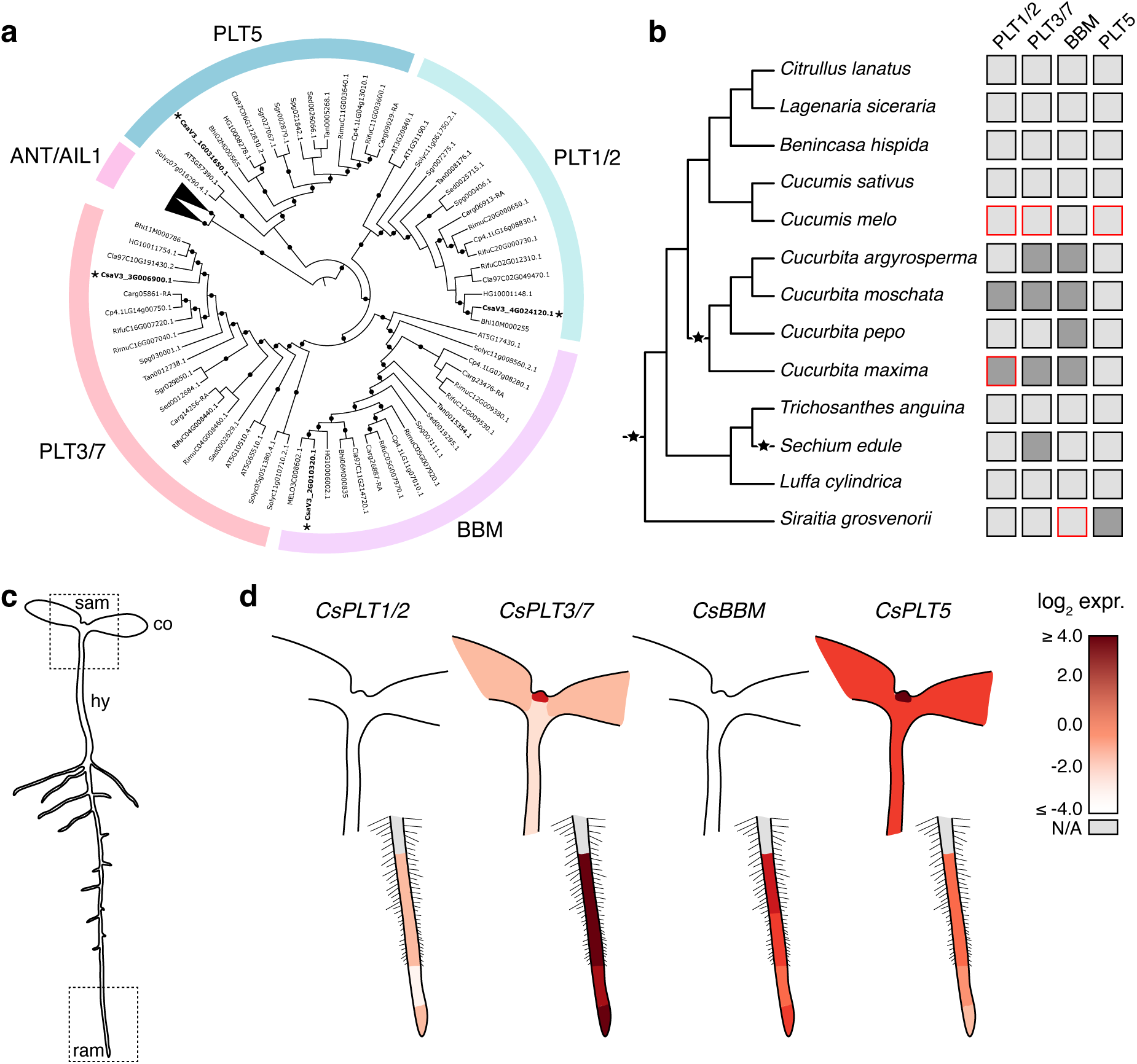
*CsPLTs* are expressed in root and shoot apices. (**a**) Maximum-likelihood phylogenetic tree of euANT transcription factors with >1 AP2 domain in 13 cucurbit species, Arabidopsis and tomato. The tree is rooted at the split between the ANT/AIL1 (collapsed) and PLT clades. Black circles correspond to bootstrap values > 70. Cucumber homologs are indicated with *. (**b**) Phylogeny and whole-genome duplication events (stars, (Ma *et al*., 2022)) in the analysed cucurbit species. Copy numbers per PLT clade per species are indicated in grey (light = 1, dark = 2). Squares with red borders indicate that the count includes one homolog with < 2 AP2 domains, inferred from synteny with the cucumber homolog (Yu *et al*., 2023), possibly because of wrong annotation. (**c**) Schematic drawing of a 6 dpg cucumber seedling. sam = shoot apical meristem, co = cotyledon, hy = hypocotyl, ram = root apical meristem. (**d**) Heatmap of cucumber seedling *CsPLT* expression in tissues demarcated by dashed rectangles in (**c**). Intensity corresponds to the log_2_ expression value (+ 1e-5).

Subsequently, we aimed to study expression patterns of these four cucumber *PLTs (CsPLT).* We germinated seeds of Chinese long inbred line ‘9930’, a wild type monoecious cucumber with a well-annotated genome (Li *et al*., 2019), and dissected various tissues at 6 days post germination (dpg). Like in Arabidopsis, *CsPLTs* were most strongly expressed in meristematic tissues (Fig. 1c,d). Whereas we detected expression of all four genes in the root tip, only *CsPLT3/7* and *CsPLT5* were expressed in shoot tissues (Fig. 1d). Transcript levels were particularly elevated in the shoot apex, containing both the shoot apical meristem and early leaf primordia. We thus inferred that only *CsPLT3/7* and *CsPLT5* were potentially involved in cucumber shoot development.

### *CsPLT3/7* and *CsPLT5* are expressed in partially overlapping domains

To increase the resolution of transcript localisation, we performed *in situ* hybridisations on longitudinal sections of 5 dpg ‘9930’ shoot apices with *CsPLT3/7* and *CsPLT5* probe sets that form a highly specific red/pink precipitate emitting fluorescence upon transcript detection. Compared to the no probe control, *CsPLT3/7* mRNA localised primarily to the summit of the SAM dome (i.e. the CZ, containing the stem cell niche), the axillary meristems, and weakly in developing leaf primordia (Fig. 2a,b,d,e). On the other hand, *CsPLT5* was expressed strongly in leaf primordia and moderately throughout the SAM, but not clearly in the *CsPLT3/7* domain (Fig. 2a,c,d,f). To verify this observation, we made cross sections through the stem cell niche and again observed that *CsPLT5* was not expressed in this region, unlike *CsPLT3/7* (Fig. 2g-l). Thus, *CsPLT3/7* and *CsPLT5* are expressed in partially overlapping expression domains and likely influence distinct processes in the SAM and leaf primordia.

**Figure 2.**
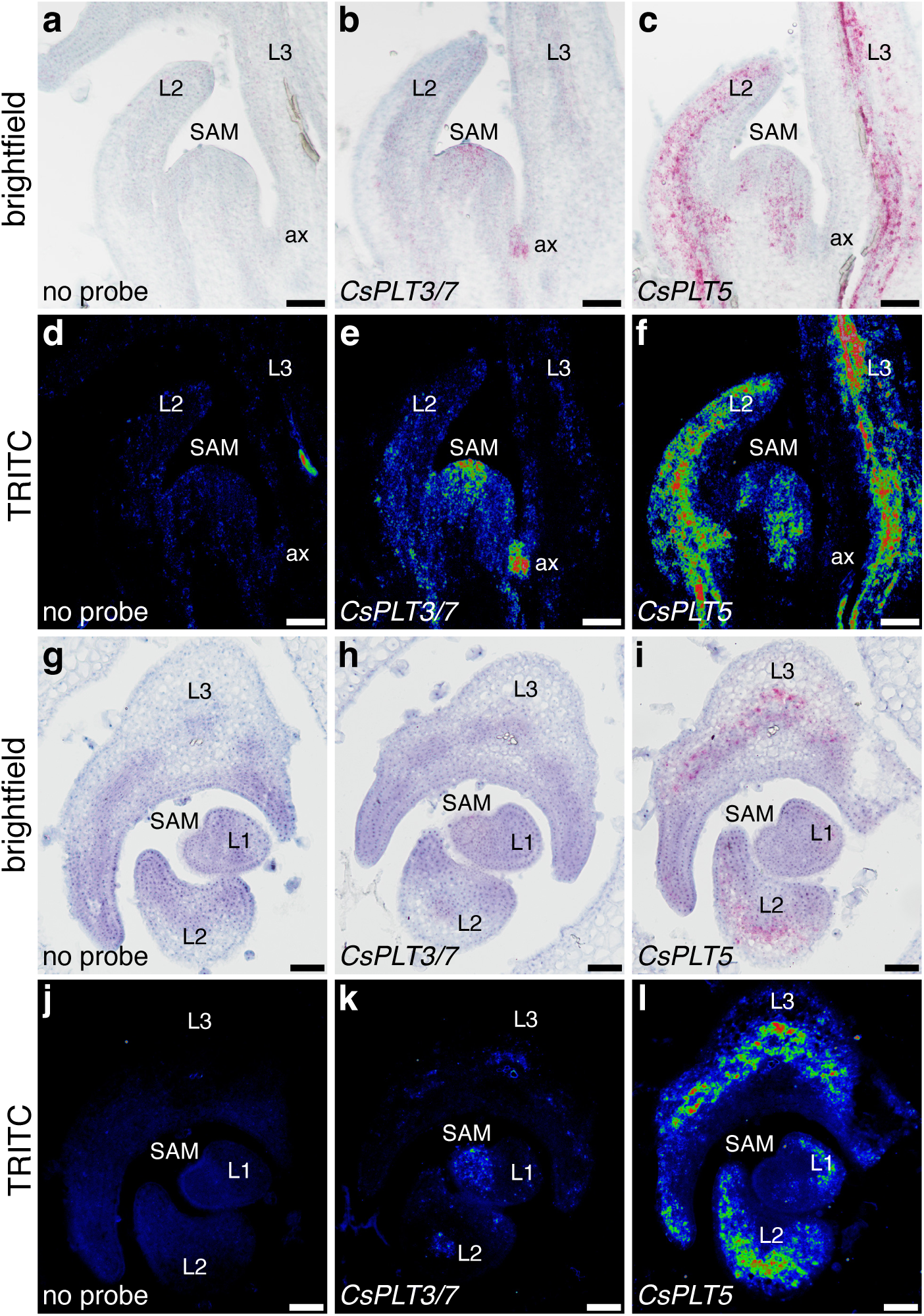
*CsPLT3/7* and *CsPLT5* are expressed the cucumber SAM. (**a-f**) Longitudinal sections of 5 dpg cucumber SAMs after *in situ* hybridisation with *CsPLT3/7* or *CsPLT5* probe sets, with a no-probe control. The brightfield images (**a-c**) correspond to the equal-exposure TRITC fluorescent images in (**d-f**). Cross sections are displayed in (**g-l**). Sections are counterstained with Gill’s Hematoxylin I. L = leaf primordium, ax = axillary meristem. Scale bars are 100 μm.

### CsPLT3/7 and CsPLT5 bind to SAM patterning genes

To gain more insight in the function of CsPLT3/7 and CsPLT5, we performed DNA Affinity Purification sequencing (DAP-seq) of genomic DNA from cucumber shoot apices. We identified 3937 and 1947 regions bound by CsPLT3/7 and CsPLT5, respectively, with strong congruence between both datasets; 1599 of 1947 (82%) CsPLT5 peak coordinates overlapped for at least 50% with a CsPLT3/7 peak (Fig. 3a, Table S1). Across two replicates, the CsPLT5 DAP seemed to have a slightly lower signal-to-noise ratio than CsPLT3/7, reflected by its smaller number of significant peaks and lower FRiP score (Table S2). Motif analysis of the summit sequences revealed that both TFs recognize a centrally enriched ANT-like consensus motif, which is very similar to that of Arabidopsis PLT3, PLT5 and PLT7 (Fig. 3b,c). The high similarity of the binding motifs, together with the extensively shared DAP peaks, suggests that CsPLTs bind the same target sequences, as is the case in Arabidopsis (Santuari *et al*., 2016; Kerstens *et al*., 2025a).

**Figure 3.**
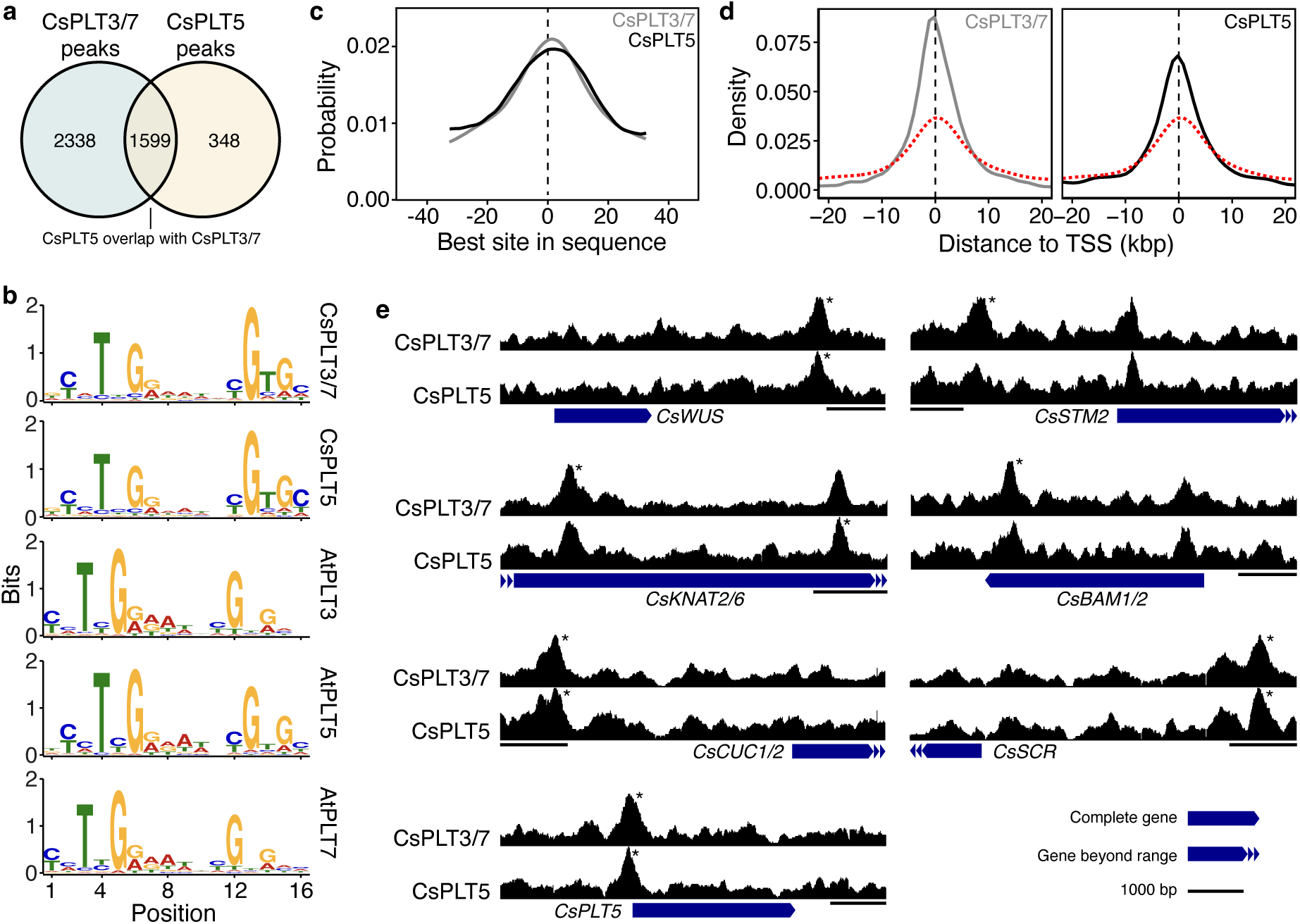
CsPLTs bind SAM-expressed genes *in vitro*. (**a**) CsPLT3/7 and CsPLT5 DAP-seq peaks identified from two replicates. Overlapping peaks are CsPLT5 peaks shared for at least 50% (of the CsPLT5 peak) with CsPLT3/7. (**b**) De novo motif discovery of CsPLT3/7 and CsPLT5, in comparison to the motifs of AtPLT3, AtPLT5 and AtPLT7 generated from inflorescence meristem DNA in Arabidopsis (Kerstens *et al*., 2025a). (**c**) Central enrichment of the motifs in (b) as calculated by CentriMo. (**d**) Distance of CsPLT DAP peaks from the nearest TSS. The red dashed line represents the by chance expected distribution of distances to the nearest TSS based on 10000 randomly selected genomic regions. (**e**) Coverage tracks of CsPLT3/7 (top) and CsPLT5 (bottom) over annotated SAM organising genes. *CsSTM2* is one of two *AtSTM* co-orthologs. * Denote significant peaks (IDR < 0.05).

We then identified CsPLT-regulated candidate genes by attributing peaks to the nearest transcription start site (TSS). Both CsPLTs showed enriched binding near TSSs over a simulated background, primarily within 5 kbp from the TSS and peaking around the central coordinate (Fig. 3d, Table S1). To narrow down relevant target genes, we considered only those genes bound by either CsPLT within a [-5 kbp, 5 kbp] range from the TSS. Additionally, we performed RNA-seq on cucumber shoot apices to select for candidate genes expressed within the SAM (Table S3). Within the curated list of 1446 candidate target genes (Table S4), we found the cucumber orthologs of *WUS* (*CsaV3_6G047050*), the Class I *KNOTTED-LIKE FROM ARABIDOPSIS THALIANA* (*KNAT*) genes *SHOOT MERISTEMLESS* (*STM*; *CsaV3_7G003300*) and *KNAT2/KNAT6* (*CsaV3_2G024800*), *CUP-SHAPED COTYLEDON1/2* (*CUC1/CUC2; CsaV3_4G033430*), *BARELY ANY MERISTEM1/2* (*BAM1/BAM2; CsaV3_6G044530*), and *SCARECROW* (*SCR; CsaV3_4G013520*), all of which are (redundantly) required for SAM formation, maintenance or size in Arabidopsis (Fig. 3e; (Barton & Poethig, 1993; Laux *et al*., 1996; Aida *et al*., 1997; DeYoung *et al*., 2006; Belles-Boix *et al*., 2006; Bahafid *et al*., 2023)). Additionally, a peak close to the *CsPLT5* TSS was observed, indicating the potential existence of an autoregulatory feedback loop (Fig. 3e). SAM PLTs thus bind to - and potentially regulate - key meristematic organizing genes in cucumber.

### CsPLT1/2 and CsPLT3/7 are required during embryogenesis

We then set out to determine the functional role of CsPLTs in shoot architecture. Using EMS mutagenesis and TILLING, we generated a library of *Csplt* mutant alleles in a gynoecious parental line (Fig. S1a; Table S5). For *CsPLT1/2*, we obtained two premature stop codon alleles at tryptophan 52 and glutamine 58, which lack both AP2 DNA-binding domains and are thus predicted to be loss-of-function alleles. For *CsPLT3/7*, we obtained the two premature stop codon alleles *Csplt3/7^W48*^* and *Csplt3/7^Q156*^*, both lacking the AP2 domains, and *Csplt3/7^R355H^*, harbouring an arginine to histidine amino acid substitution within the second AP2 domain. We also obtained the *Csbbm^W317*^*, *Csplt5^R289*^* and *Csplt5^L189F^*alleles, in which the proteins are truncated at the end of the first or second AP2 domain or harbouring a leucine to phenylalanine substitution in the first AP2 domain, respectively.

During propagation of the TILLING mutants, we obtained lower frequencies of mutants than expected, suggesting a role for CsPLTs during embryogenesis. Indeed, homozygous Cs*plt3/7^W48*^* and *Csplt3/7^Q156*^* mutants occurred at significantly lower frequencies in the segregating offspring from a selfed heterozygous plant than heterozygous or wild type genotypes (Fig. S1b), which we did not observe for any of the other genotypes in this study. To uncover whether specific combinations of *Csplt* alleles are fully embryonic lethal, we generated various *Csplt* double mutant combinations through crossing. Segregation of single *Csplt* alleles in a wild type background in this experiment again demonstrated that only *Csplt3/7^Q156*^* mutants were underrepresented in the progeny (Fig. 4a). Strikingly, we did not find a single homozygous *Csplt3/7^Q156*^*seedling in 337 plants with a homozygous *Csplt1/2^W52*^* mutation, nor a single homozygous *Csplt1/2^W52*^* seedling in 139 plants homozygous for *Csplt3/7^Q156*^* (Fig. 4b). Moreover, out of all double mutants, only the frequencies of *Csplt3/7^Q156*^* in the *Csplt1/2^W52*^*background and vice versa were significantly reduced in comparison to the respective single mutant allele frequencies (Fig. 4b). The occurrence of the homozygous *Csbbm^W317*^* genotype in the *Csplt3/7^Q156*^* background and vice versa also seemed to be slightly underrepresented, albeit not significantly (Fig. 4b). Corroborating these findings, we found that expression of *GUS* from the *CsPLT1/2* and *CsPLT3/7* promoters in Arabidopsis occurred during early embryogenesis (Fig. S2). We conclude that the combined loss of CsPLT1/2 and CsPLT3/7 is embryonic lethal.

**Figure 4.**
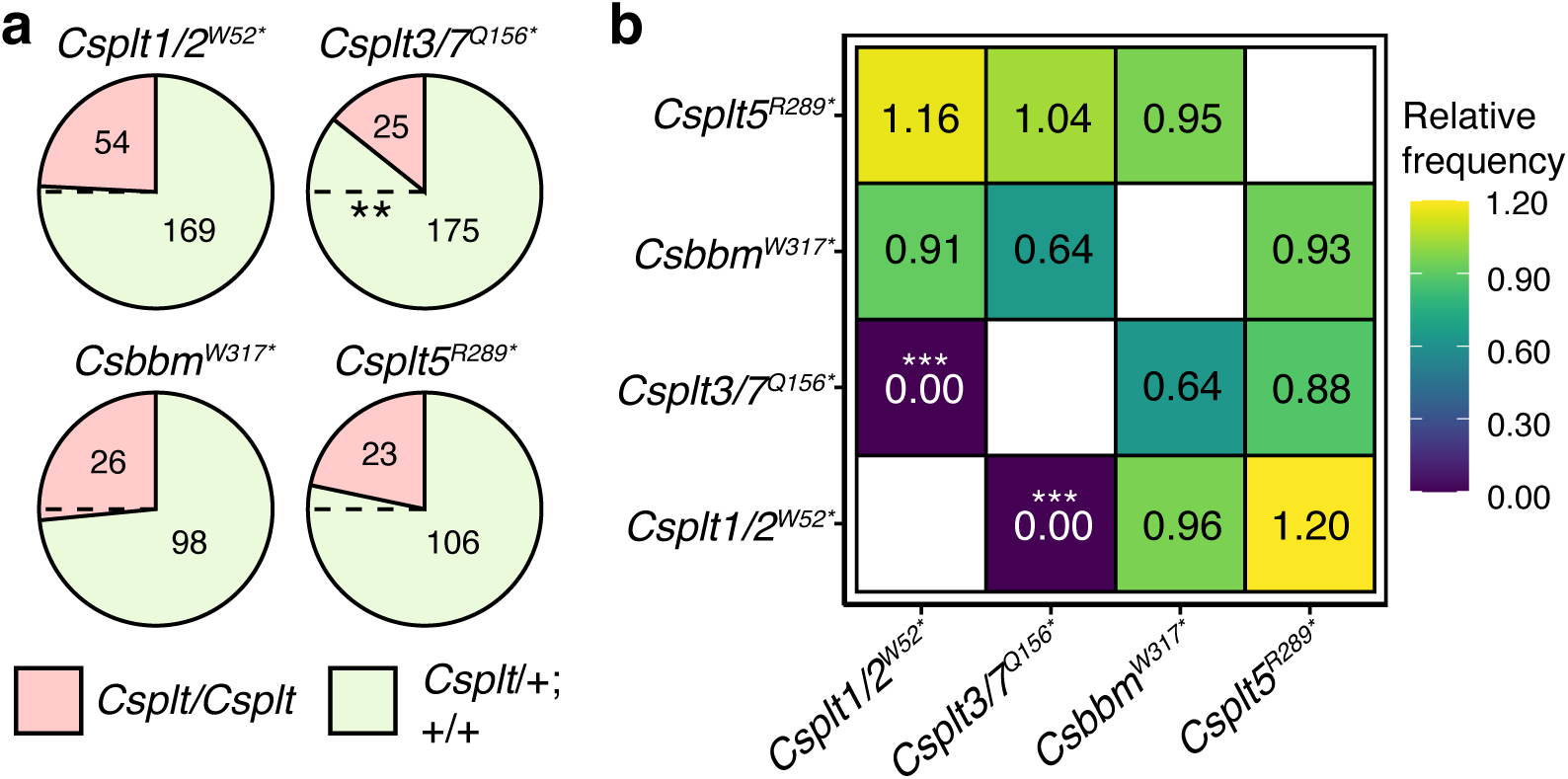
*Csplt1/2^W52*^ Csplt3/7^Q156*^* double mutants are embryonic lethal. (**a**) Pie charts displaying the proportion of homozygous mutants (red) and heterozygous + wild type seedlings (green) per analysed TILLING allele, with counts indicated. The dashed line indicates the expected frequency of 0.25 based on Mendelian segregation. (**b**) Heat map matrix showing homozygous *Csplt* genotype frequency in different homozygous *Csplt* backgrounds, relative to the frequency of the respective single mutant genotype in (**a**). For example, the upper left square is the occurrence frequency of the homozygous *Csplt5^R289*^* genotype in a homozygous *Csplt1/2^W52*^* background, relative to the frequency of the homozygous *Csplt5^R289*^*single mutant. Self-comparisons are plotted in white. Statistics are Benjamini-Hochberg (BH) and Yates(Y)-corrected z-tests (freq. < 0.25 (**a**) or freq. < freq(homozygous single mutant) (**b**); ** p = 0.003; *** p = 3.8e-4).

### *Csplt3/7* mutants lack a functional SAM at germination

Given the role of *CsPLT3/7* during embryogenesis and its expression in the stem cell niche, we next asked if shoot development in young seedlings with a loss-of-function *Csplt3/7* allele (i.e. *Csplt3/7^Q156*^*) was affected. In mature embryos dissected from wild type seeds, a single leaf primordium was visible (Fig. 5a). At 6 dpg, this primordium grew into a distinct first true leaf (L1), which after further dissection revealed a much smaller second leaf (L2) and recently initiated primordium (L3), with a dome-shaped SAM at the centre (Fig. 5b-d). Contrarily, *Csplt3/7* embryos already had two primordia before germination, which developed into two equally sized leaves oriented at 180° from each other at 6 dpg (Fig. 5e-g). Strikingly, the two primordia were separated by a flat apex bearing no resemblance to a SAM (Fig. 5h). Microscopic analysis revealed the presence of trichomes across the entirety of the flat apex - a hallmark of differentiation, which in wild types occurred only in the tissue surrounding the SAM (Fig. 5i,j). In 15 dpg *Csplt3/7* seedlings, numerous leaf-like structures emerged from the from the apex, at which point wild type plants possessed one large and one small leaf (Fig. 5o-n). This indicates that the *Csplt3/7* apex retains meristematic properties despite the initial absence of a functional SAM. The *Csplt3/7* leaves displayed roughly synchronous development at later time points, eventually establishing new functional secondary meristems at leaf junctions in 29 of 30 plants at 45 dpg (Fig. 5p,q). We conclude that CsPLT3/7 is required for embryonic SAM establishment or maintenance during the formation of the first leaf primordia.

**Figure 5.**
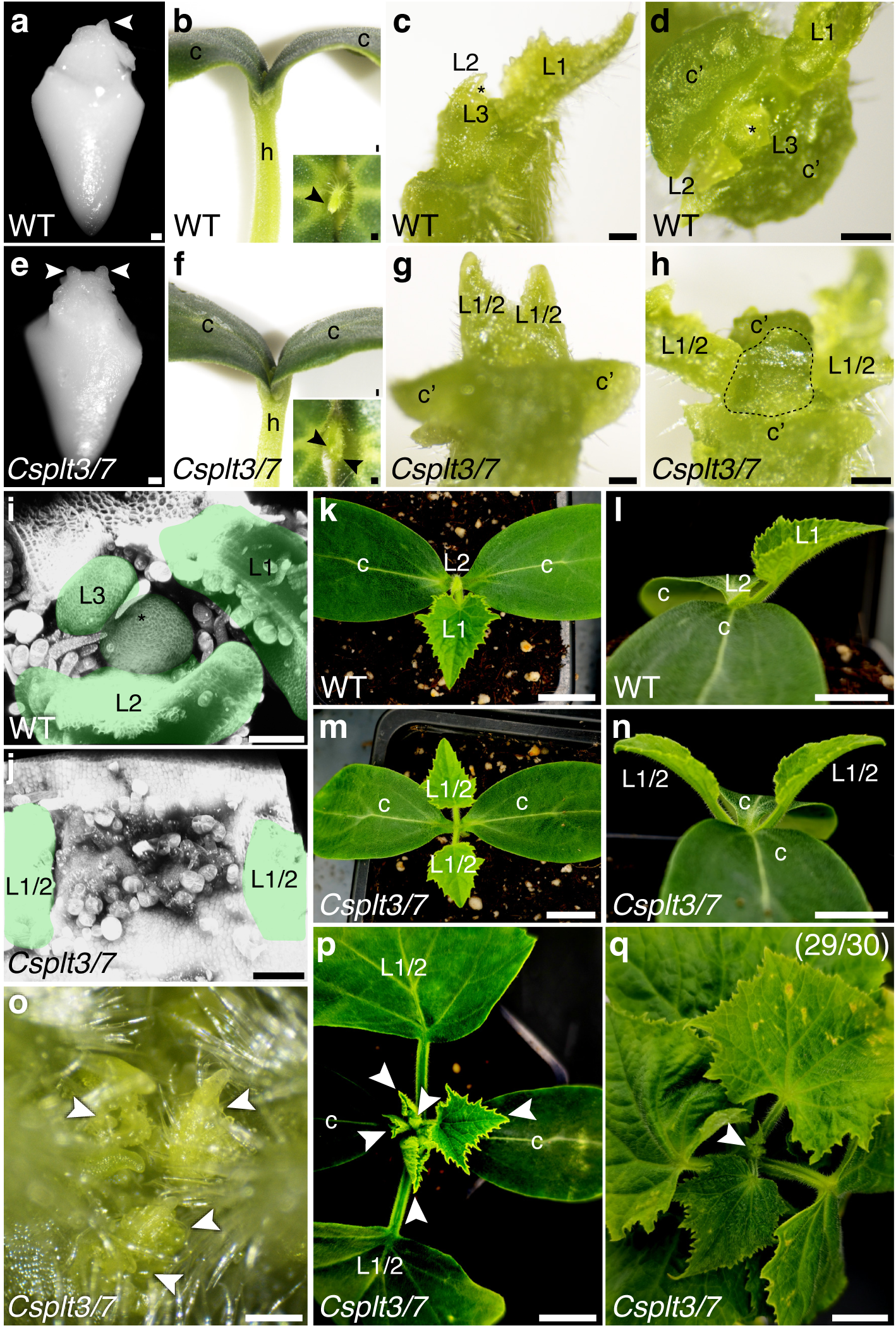
CsPLT3/7 regulates leaf initiation in seedlings. (**a**) 4 hours past imbibition WT embryo, (**b**) 6 dpg seedling with top-view inset (**b**), shoot apex closeup side view (**c**) and top view (**d**) compared to same-age *Csplt3/7^Q156*^*(**e**-**h**). Arrowheads in point to initiated leaves. The dotted lines indicate the site where a SAM would have been expected. (**i,j**) Maximum projections of shoot apices, with leaves (scars) indicated in yellow. The SAM is coloured green. Cell wall staining is SR2200. (**k**-**n**) 15 dpg seedlings, with a closeup (**o)** showing de novo leaf formation in *Csplt3/7^Q156*^* (arrowheads). (**p**) 22 dpg *Csplt3/7^Q156*^* plant with newly formed leaves (arrowheads) and appearance of a new SAM at 45 dpg (**q**, arrowhead). Abbreviations: h = hypocotyl, c = cotyledon, c’ = resected cotyledon, Lx = leaf (order of initiation), * = SAM. Scale bars are 200 μm in (**a**-**h**), 100 μm in (**i**,**j**), 75 μm in (**o**) and 1 cm in (**k**-**n**, **p, q**).

### CsPLT3/7 shapes mature shoot architecture

We then grew all obtained TILLING mutant lines, including *Csplt3/7^Q156*^*, in the greenhouse in order to evaluate shoot architecture traits in mature plants. At 6 weeks after planting, homozygous *Csplt1/2^W52*^*, *Csplt1/2^Q58*^*and *Csbbm^W317*^* mutants did not display shoot phenotypes and had a similar internode length compared to wild type plants, in line with absence of expression in the SAM (Fig. S3). Likewise, *Csplt5^L189F^*and *Csplt5^R289*^* mutants exhibited no conspicuous phenotypes, although internodes in the latter line were slightly (4%; 9.3 vs. 9.7 cm) shorter than the wild type (Fig. S3). To the contrary, the three *Csplt3/7* mutants each exhibited clear defects in shoot development. In all three lines we observed aberrant internode length, ranging from near absence to lengths up to 20 cm, in contrast to highly regular lengths between 5 and 10 cm in wild type plants (Fig. 6a-g). Occasionally, *Csplt3/7^Q156*^* plants formed temporary stretches of extremely short internodes succeeded by extremely long ones, resulting in pairs of leaves separated by long stem sections (Fig. 6c). Since these stem arrangements occurred spontaneously amidst medium-sized internodes, we wondered if certain internode sizes occurred preferentially in succession. We therefore used a Markov chain framework to determine the probabilistic transitions between three internode length classes in *Csplt3/7^Q156*^*, i.e. short, medium-length, and long internodes. Whereas medium-length internodes - constituting the majority of measured internodes (Fig. S4a) - were most frequently (81%) succeeded by medium-length internodes, short internodes were preferentially (62%) followed by long internodes, and long internodes were succeeded equally likely (48% each) by short internodes or medium-sized internodes (Fig. S4b). Short internodes occurred sporadically (19%) in succession, but this only happened very rarely (3%) for long internodes (Fig. S4). These data suggest that after an aberrant internode length is generated, subsequent internodes generally remain extreme in size.

**Figure 6.**
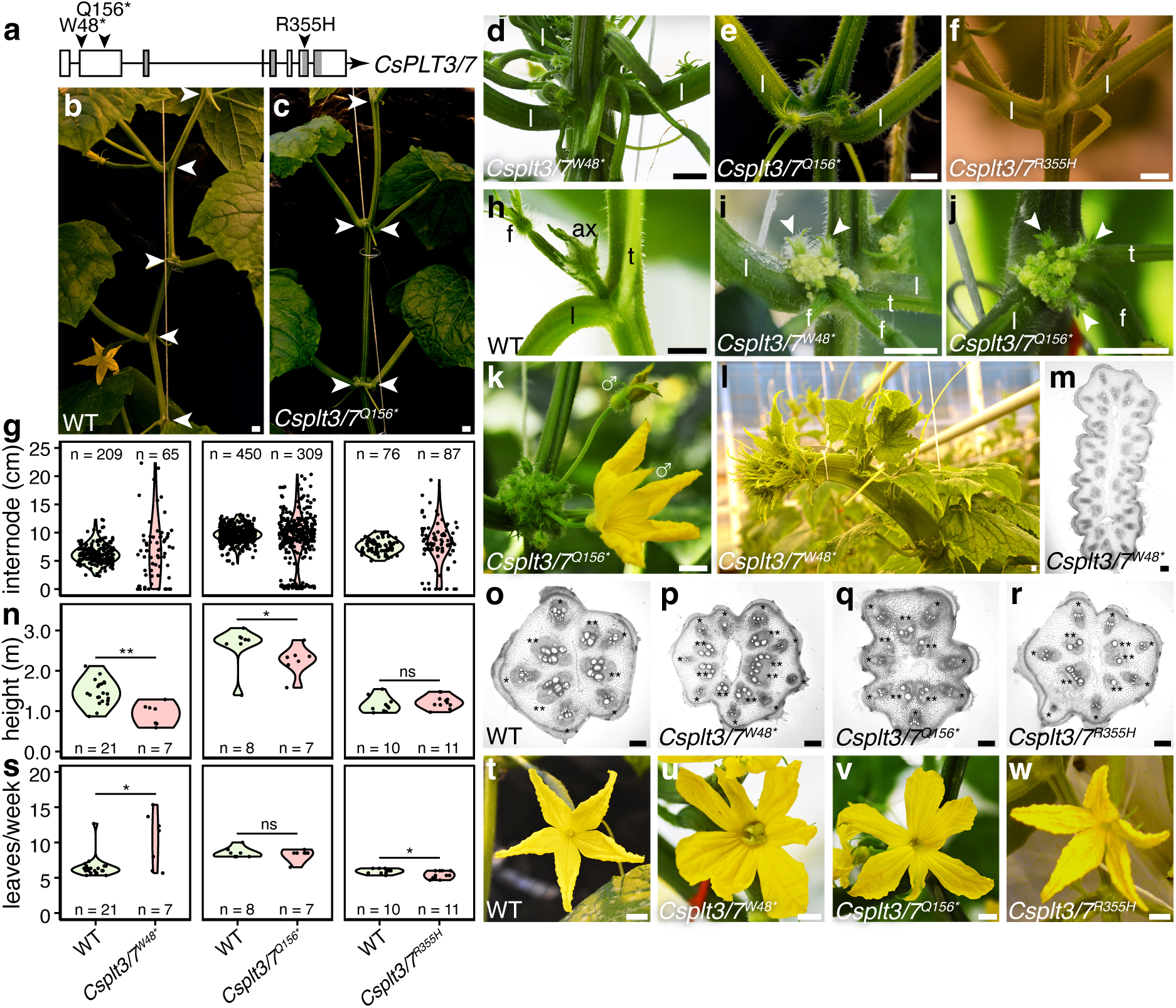
CsPLT3/7 regulates shoot architecture and flower development. (**a**) Gene model displaying *Csplt3/7* alleles. AP2 domains are indicated in grey. (**b**-**c**) 6-week-old WT or *Csplt3/7^Q156*^* shoots, after removal of lower axillary shoots and tendrils. Arrowheads point towards nodes. (**d**-**f**) Near-co-initiated nodes of *Csplt3/7^W48*^*, *Csplt3/7^Q156*^* and *Csplt3/7^R355H^*. (**g**) Internode length of the three alleles, with each dot representing individual internodes. Note that fasciated regions of *Csplt3/7^W48*^* were not counted. (**h**-**j**) Axil morphology of WT, *Csplt3/7^W48*^* and *Csplt3/7^Q156*^* plants. Arrowheads point to initiated flowers from the callus-like tissue. (**k**) Older axil of *Csplt3/7^Q156*^*showing extreme flower production and two male flowers. (**l**) Fasciation in *Csplt3/7^W48*^* and a cross section through the stem (**m**). (**n**) Shoot height of WT and *Csplt3/7* mutants. Statistics are two-tailed Wilcoxon rank-sum tests (left to right: p = 4.1e-4, p = 0.03, p = 0.26). (**o**-**r**) Stem cross sections of WT and *Csplt3/7* lines. * and ** denote outer and inner vascular bundles, respectively. (**s**) Average leaf formation per week of WT and *Csplt3/7* mutants. Statistics are two-tailed Wilcoxon rank-sum tests (left to right: p = 0.03, p = 0.71, p = 0.03). (**t**-**w**) Flower morphology of WT and *Csplt3/7* lines. Scale bars in (**b**-**f**, **h**-**l**, **t**-**w**) are 1 cm, and 0.1 mm in (**m**, **o**-**r**). Abbreviations: l = leaf, ax = axillary shoot, f = flower, t = tendril.

We observed that in the two presumed loss-of-function mutants, *Csplt3/7^W48*^* and *Csplt3/7^Q156*^*, callus-like growths consistently emerged from the axils, from which large numbers of female and sometimes male flowers emerged later, despite the background line being gynoecious and normally producing only 1 female flower per node (Fig. 6h-k). While axillary shoots could still form, their emergence appeared inconsistent (Fig. 6h-j). In wild type plants, the first axillary shoots generally developed from the axils of the 8^th^ to 10^th^ node (from the shoot apex) and by the 14^th^ node were consistently present (Fig. S5). Conversely, in same-age *Csplt3/7^Q156*^* plants, axillary shoots were routinely absent and were formed infrequently (Fig. S5). Such shoots also exhibited the internode phenotype of the main shoot (Fig. S6). Tendril development appeared unaffected. We also observed that *Csplt3/7^W48*^*stems often (10/17) underwent fasciation during growth (Fig. 6l,m), which never occurred in *Csplt3/7^Q156*^*and *Csplt3/7^R355H^*. Furthermore, shoot height of the loss-of-function mutants was reduced (Fig. 6n), and leaf production - a proxy for plastochron - was increased in the fasciating *Csplt3/7^W48*^*line, but marginally decreased in *Csplt3/7^R355H^* (Fig. 6s). Since *Csplt3/7^W48*^*mutants experienced fasciation, we wondered whether stem architecture of *Csplt3/7^Q156*^*and *Csplt3/7^R355H^* was more subtly affected. In cross sections of internodes of *Csplt3/7^Q156*^* and *Csplt3/7^R355H^*mutants, as well as in non-fasciating *Csplt3/7^W48*^* stems, supernumerary ridges and vascular bundles were present compared to 5-ridged wild type stems with 4 inner and 5 outer vascular bundles (Fig. 6o-r). Finally, flower morphology of the two presumed loss-of-function mutants was disturbed, exhibiting alternative sepal/petal numbers and shape (Fig. 6t-w). We measured this phenotype in more detail in *Csplt3/7^Q156*^*. Whereas wild type flowers (n = 120) invariably formed 5 sepals, 5 petals and 3 carpels, *Csplt3/7^Q156*^*flowers developed a wider range (between 3 and 9) of sepals and petals (Fig. S7). Moreover, we observed partial sepal-sepal or sepal-petal fusions in ∼40% (13/32) of flowers, and ∼6% (2/32) of flowers exhibited formation of 4 carpels instead of 3 (Fig. S7). We conclude that CsPLT3/7 regulates internode length, axillary shoot and flower emergence, and floral morphology in mature plants.

### CsPLT3/7 regulates phyllotaxis

Since loss of Arabidopsis SAM- and IM-expressed PLTs, i.e. PLT3, PLT5 and PLT7, reduces robustness of the phyllotactic spiral (Prasad *et al*., 2011; Pinon *et al*., 2013; Kerstens *et al*., 2025a), we asked if *Csplt3/7* mutants experienced more irregular circumferential leaf patterning. Using a custom-made hinged protractor (Robertson *et al*., 2025), we manually quantified divergence angles in 22.5° bins between successive leaves in *Csplt3/7^W48*^*, *Csplt3/7^Q156*^* and *Csplt3/7^R355H^*, and the respective wild type sister progeny per line. Unlike Arabidopsis, in which spiral phyllotaxis is characterised by pattern convergence around the ‘golden angle’ (∼137.5°; within a 135±22.5° angle bin), the most frequently occurring divergence angle bin in all three wild type cucumber lines was 157.5±22.5° (Fig. 7a). These data suggest that spiral phyllotaxis does not typically conform to the golden angle on mature cucumber stems. In *Csplt3/7^W48*^*, in which we could only analyse a handful of divergence angles from non-fasciating stems with a clear initiation order, divergence angle distribution was much broader than in its wild type sister progeny, suggesting reduced robustness of the phyllotactic spiral (Fig. 7a). On the other hand, in both *Csplt3/7^Q156*^* and *Csplt3/7^R355H^* the overall patterning regularity seemed unaffected. We instead found a significant shift towards a distichous pattern, in which leaves are consecutively positioned at 180±22.5° angles from each other (Fig. 7a,b). The distichous pattern only manifested occasionally; a large proportion of divergence angles also approached 157.5±22.5°.

**Figure 7.**
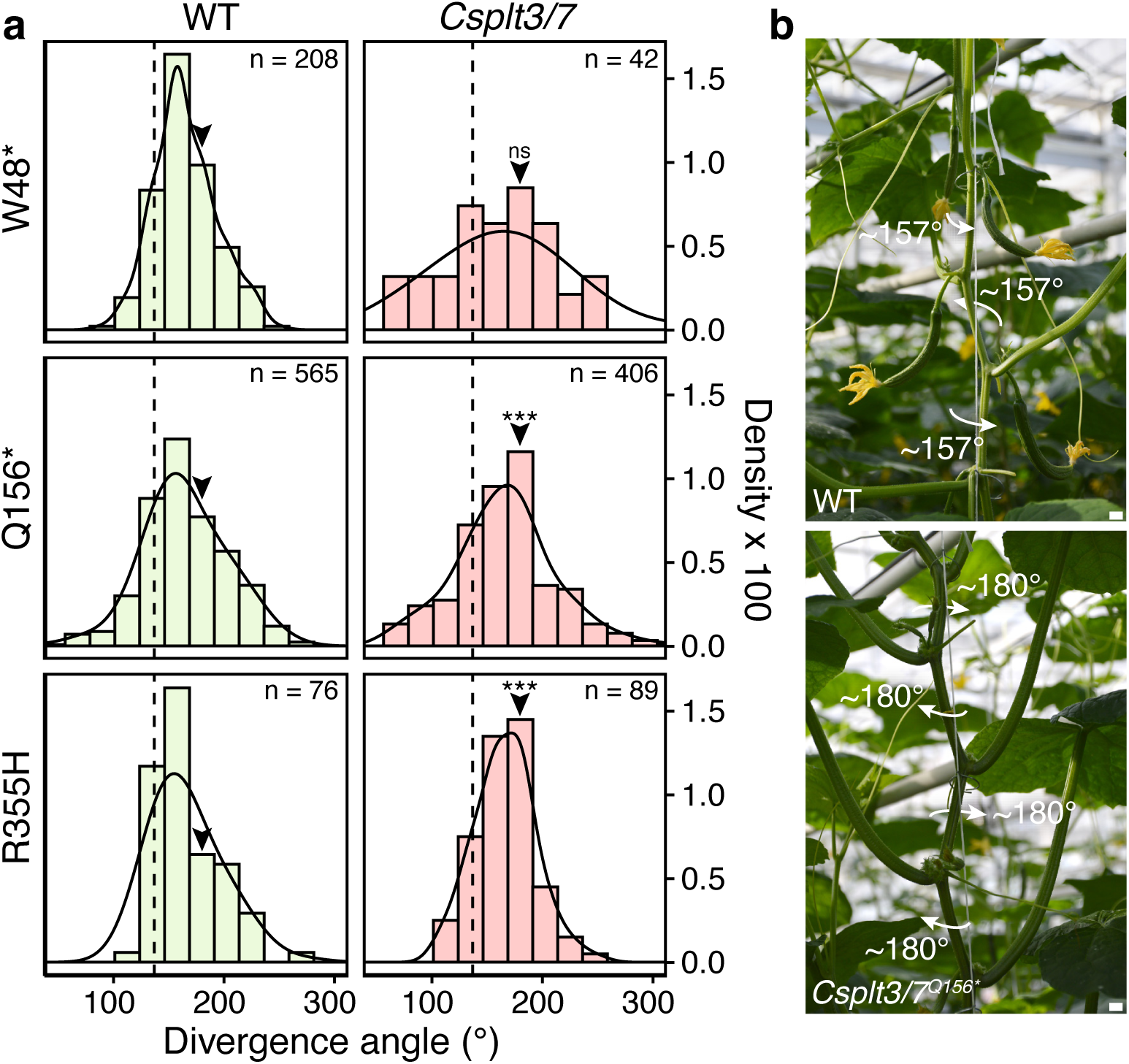
*Csplt3/7* mutants exhibit altered phyllotaxis. (**a**) Histograms and density plots displaying the occurrence of divergence angles within 22.5° bins. The dashed line indicates the golden angle (137.5°), and arrowhead points towards the bin containing 180°. Significance from BHY-corrected pairwise z-tests (top to bottom: p = 0.71, p = 5.0e-5, p = 5-4e-4). (**b**) Example sections of cucumber stems displaying “spiral” (upper; counterclockwise) and distichous (lower; clockwise) phyllotactic patterning. Scale bar is 1 cm.

Given that *Csplt3/7^Q156*^* mutants formed extreme internodes and we previously showed that Arabidopsis phyllotaxis is strongly chirality-dependently modified by stem torsion acting on internode length (Kerstens *et al*., 2025a), we investigated if its shift towards distichous phyllotaxis is caused by stem torsion. We first quantified the degree to which torsion occurred and affected wild type phyllotaxis in our growth conditions. Unlike Arabidopsis, in which stems twisted persistently in a counterclockwise manner (Kerstens *et al*., 2025a), cucumber stems twisted almost equally in clockwise and counterclockwise directions with on average a minute preference to clockwise twisting (μ = -0.17°; Fig. S8a,b). Using a trigonometric representation of the stem that could previously predict the effect of torsion on phyllotaxis (Landrein et al., 2013), we predicted that in an average internode with an average stem radius, the divergence angle would be displaced 7° in the clockwise direction (Fig. S8c,d; Equation 1). We next plotted divergence angles against internode length separately for clockwise and counterclockwise-turning meristems. Divergence angles of clockwise and counterclockwise-turning meristems increased and decreased with longer internodes, albeit insignificantly, as a consequence of (on average) slight clockwise stem torsion (Fig. S8e). Accordingly, the average divergence angle of clockwise-turning meristems was slightly larger than that of counterclockwise-turning meristems (Fig. S8f). When we repeated this analysis for *Csplt3/7^Q156*^* and its wild type sister progeny in a different growth season, however, the relationship between internode length and divergence angle appeared inverted and was again not significant (Fig. S8g). Moreover, in internodes of equal length (9-11 cm), the proportion of 180° angles in the mutant was still enriched (Fig. S8h). We conclude that the phyllotaxis defect of *Csplt3/7^Q156*^*, and likely per extension, that of *Csplt3/7^R355H^*, cannot be explained by a cooperative effect of stem torsion and altered internode length. Thus, loss of functional CsPLT3/7 shifts phyllotaxis to larger divergence angles.

## Discussion

In this study, we have shown that *PLT* genes in cucumber are expressed in meristematic tissues, with *CsPLT3/7* and *CsPLT5* being the only two detected in the SAM (Fig. 1). Within the SAM, both TFs recognised the same consensus motif and bound to SAM organising genes, but only *CsPLT3/7* was expressed in the stem cell niche (Fig. 2,3). Lack of CsPLT3/7 distorted embryogenesis (Fig. 4), as well as embryonic SAM establishment or maintenance (Fig. 5). Mature *Csplt3/7* shoots occasionally displayed irregular internode lengths and axillary organ development and shifted phyllotactic patterning (Fig. 6,7). Altogether, we firmly establish CsPLT3/7 as a major regulator of cucumber shoot development and architecture.

Despite the use of TILLING lines, several lines of evidence support that the observed phenotypes are caused by mutations in *CsPLT3/7*. Lack of initial SAMs, leaf initiation and axil defects in *Csplt3/7^W48*^* and *Csplt3/7^Q156*^* are in congruence with *CsPLT3/7* expression domains based on our *in situ* hybridizations. The fact that internode length and stem architecture aberrancies occurred in three independent TILLING alleles, and the same axil and flower defects occurred in the premature stop alleles *Csplt3/7^W48*^* and *Csplt3/7^Q156*^*, reinforces the validity of *CsPLT3/7* being the causal gene. Why only *Csplt3/7^W48*^*mutant stems fasciate despite both stop alleles truncating the transcript in the first exon, remains unclear; perhaps another TILLING lesion in close proximity to *CsPLT3/7* co-segregated with *Csplt3/7^W48*^*, further exacerbating the phenotype. Notably, a similar phenotype has been described for an amino acid substitution allele of *Csclv1*, in which stem fasciation occurred and flowers exhibited supernumerary petals and carpels (Cheng *et al*., 2022). Supernumerary floral organs, but not fasciation, was also described for *Csclv3* mutants and RNAi lines (Che *et al*., 2020; Han *et al*., 2024). We did not identify notable shoot phenotypes for other *Csplt* mutants, which was expected for the non-SAM-expressed *CsPLT1/2* and *CsBBM*, but not *CsPLT5*. Strictly, we cannot exclude that *CsPLT5* plays an additional role in the shaping shoot architecture due to the nature of the TILLING alleles, i.e. the weak *Csplt5^L189F^* and the *Csplt5^R289*^* alleles, which lacks only the most terminal portion of the second AP2 domain. While targeted gene editing strategies such as CRISPR/Cas9 would be required to resolve such ambiguities, stable transformation of the recalcitrant cucumber remains a persistent bottleneck that only select laboratories have been able to overcome.

Even though *Csplt3/7* mutant shoots resumed indeterminate growth, the regularity of wild type internode length was lost. The mechanism behind the formation of extremely short and long internodes remains a matter of speculation at this point. One possible scenario is that *Csplt3/7* SAMs are unable to rhythmically develop leaf primordia due to disruption of auxin signalling and transport. Short internodes would then result from the near simultaneous formation of two primordia, for example through increased meristem size (Landrein *et al*., 2015) or permissive inhibitory fields (Besnard *et al*., 2014). Extremely long internodes could arise from the inability to establish auxin maxima, as we previously observed in inflorescences of the Arabidopsis *plt3 plt5 plt7 pin1^T600I^* quadruple mutant (Kerstens *et al*., 2025a). Our observations that extreme internodes preferentially occur in succession and that phyllotactic patterns are shifted, lends additional support to the scenario that the internode phenotype results from primordium initiation defects. Importantly, this would fit with the proposed hypothesis that shoot meristematic PLTs confer patterning robustness to the SAM (Kerstens *et al*., 2025a), acting within the CZ (Pinon *et al*., 2013), in which *CsPLT3/7* was expressed. Nevertheless, we strictly cannot exclude that extreme internodes result instead from variable inclusion of cells in internodes due to increased and decreased activity of the rib meristem.

Translating findings from the well-studied Arabidopsis to more distantly related species is a contemporary challenge with great impact on application potential. The nature of PLT TFs, both in regard to their conserved phylogenomic traits and functions throughout development, makes them attractive candidates to put translational hypotheses to the test. Our study shows that PLTs control meristematic processes in cucumber, as in Arabidopsis. On a whole-tissue scale, we additionally observed involvement of the same PLT clades on the root-shoot axis: *CsPLT1/2* and *CsBBM* exhibited root-specific expression, while *CsPLT3/7* and *CsPLT5* were also expressed in the shoot apex, which conforms to the expression patterns of orthologous *AtPLTs* (Fig. 1d) (Galinha *et al*., 2007; Prasad *et al*., 2011). Nevertheless, we observed clear distinctions between Arabidopsis and cucumber. Within the shoot apex, *CsPLT3/7* and *CsPLT5* were expressed in partially overlapping domains, with only *CsPLT3/7* transcripts localising to the stem cell niche. This contrasts with Arabidopsis, in which *AtPLT3*, *AtPLT5* and *AtPLT7* are all expressed in this zone (Prasad *et al*., 2011). Additionally, the observed expression of *CsPLT3/7* in axils was never reported for any *PLT* in Arabidopsis, perhaps reflecting its different growth habit or lower complexity compared to the cucumber axil. Distinctions extend to the developmental processes CsPLTs are involved in. *Csplt1/2 Csplt3/7* double mutants proved embryonic lethal, whereas in Arabidopsis, embryo lethality is only exhibited by *plt2 bbm* mutants (Chen *et al*., 2022; Kerstens *et al*., 2024). In the related cucurbit watermelon (*Citrullus lanatus*), homozygous seeds of *Clplt1/2 Clbbm* mutants could also be recovered (Liu *et al*., 2025). It was previously described that Brassicaceae *BBM* orthologs are positioned in a distinctive synteny context different from other eudicots (Kerstens *et al*., 2020), which might indicate that zygotic expression of *BBM* is not ancestral and might be controlled by different PLTs in other lineages, for instance in Cucurbitaceae.

Another distinction is that we observed a broad requirement for *CsPLT3/7* during seedling and shoot development, which is much more extreme than in Arabidopsis *Atplt3 Atplt7* (Kerstens *et al*., 2025a). In young *Csplt3/7^Q156*^* seedlings, leaves emerge de novo from the tissue where the SAM should have formed, which harbours striking similarities to *wus* mutants in Arabidopsis and Petunia (*Petunia hybrida*) (Laux *et al*., 1996; Stuurman *et al*., 2002). Mature *Atwus* embryos and *Phwus* seedlings lack SAMs, possess a flat apex, and eventually produce multiple leaf primordia and secondary SAMs across the apex in a stop-and-grow fashion (Laux *et al*., 1996; Stuurman *et al*., 2002). Secondary *Csplt3/7^Q156*^* SAMs did not terminate, however, suggesting that CsPLT3/7 is only required in the establishment or maintenance of the embryonic SAM. Since we observed binding of CsPLT3/7 near *CsWUS*, we hypothesise that the phenotype results from misregulation of *CsWUS* during early development. Although the *AtPLT3* and *AtWUS* genetic pathways interact in Arabidopsis (Mudunkothge & Krizek, 2012), SAM termination as a consequence of PLT absence does not occur in this species (Prasad *et al*., 2011). This suggests that the regulation of SAM cell fate in cucumber is wired differently. In line with this, *CsCLV3* is specifically expressed underneath the stem cell niche together with *CsWUS* (Che *et al*., 2020), which is incompatible with the canonical CLV3-WUS negative feedback loop in stem cell maintenance. Although perhaps another CsCLE family member could fulfil the function of CLV3, it is also possible that CsPLTs are responsible for stem cell maintenance, as is the case in the Arabidopsis root apical meristem (Aida *et al*., 2004). Another example that advocates differential SAM wiring in cucumber, is that the TF LEAFY, which localises to floral meristems in Arabidopsis (Weigel *et al*., 1992), is expressed in the cucumber SAM and is required for SAM maintenance in a potentially CsWUS-dependent manner (Zhao *et al*., 2018). A broader requirement for PLT3/7 TFs than in Arabidopsis was also recently reported in tomato inflorescences. Lack of SlPLT3 and SlPLT7 caused extreme branching through the disturbed regulation of *SEPALLATA* homologs (Zebell *et al*., 2025). Moreover, these inflorescences exhibited meristem overproliferation (Zebell *et al*., 2025), which we also observed in cucumber axils (Fig. 6i,j). Altogether, our findings highlight that the processes controlled by orthologous genes can shift across species, even in functionally redundant gene families such as *PLTs*. It thus becomes evident that our understanding of SAM regulatory networks across angiosperms, including cucumber, remains limited.

## Methods

### Plant material and gene identifiers

’Chinese long 9930’ and ‘long’-type parental inbred cucumber plants were propagated at Rijk Zwaan Breeding B.V. The generated mutants in the parental inbred line background and wild type sister progeny were analysed in the following generations: *Csplt1/2^W52*^*(F_2_BC_3_), *Csplt1/2^Q58*^* (F_2_BC_3_), *Csplt3/7^W48*^* (F_2_ from crossed M_2_), *Csplt3/7^Q156*^*(mature plants: M_2_ and F_2_BC_3_; seedlings: F_3_BC_4_), *Csplt3/7^R355H^*(M_2_), *Csbbm^W317*^* (F_2_BC_3_), *Csplt5^L189F^*(M_2_), *Csplt5^R289*^* (F_2_BC_3_). Double mutant genotyping was performed on F_2_ progeny obtained from two independent lines, segregating either for *Csplt1/2^W52*^, Csplt3/7^Q156*^* and *Csbbm^W317*^*, or for *Csplt1/2^W52*^*, *Csplt3/7^Q156*^*and *Csplt5^R289*^*. These segregating lines were obtained through repeated crossing of fixed F_3_BC_3_ lines. The F_2_ progeny segregating for *Csbbm^W317*^*and *Csplt5^R289*^* was obtained through crossing of M_2_ lines. *CsPLT1/2* is *CsaV3_4G024120*, *CsPLT3/7* is *CsaV3_3G006900*, *CsBBM* is *CsaV3_2G010320*, and *CsPLT5* is *CsaV3_1G031650*. Arabidopsis plants were of ecotype Col-0.

### Phylogenetics

Representative PLT protein sequences of *Citrullus lanatus* (97103 v2.5), *Lagenaria siceraria* (Hangzhou Gourd v1), *Benincasa hispida* (B227 v1), cucumber (Chinese long v3 cv 9930), *Cucumis melo* (DHL92 v4), *Cucurbita argyrosperma* (SMH-JMG-627 v2), *Cucurbita moschata* (Rifu v1), *Cucurbita pepo* (MU-CU-16 v4.1), *Cucurbita maxima* (Rimu v1.1), *Trichosanthes anguina* (v1), *Sechium edule* (v1), *Luffa cylindrica* (v1) and *Siraitia grosvenorii* (Qingpiguo v1) were retrieved from CuGenDBv2 (Yu *et al*., 2023). Arabidopsis (Araport 11) and tomato (ITAG4.0) sequences were obtained from PLAZA 5.0 (Van Bel *et al*., 2022). Proteomes were screened for sequences with > 1 AP2 domain using HMMER v3.2.1 (hmmer.org), then aligned with MAFFT v7 (Katoh *et al*., 2017), using a BLOSUM80 substitution matrix and FFT-NS-i strategy, then trimmed with trimAl v1.2rev59 (Capella-Gutiérrez *et al*., 2009), using the ‘-gt 0.5’ and ‘-cons 0.7’ flags. The trimmed sequences were aligned with IQ-TREE v1.6.10 with 1000 ultrafast bootstraps (Nguyen *et al*., 2015) and visualised with iTOL v6.8.2 (Letunic & Bork, 2019). PLT clades were defined based on the presence of Arabidopsis orthologs.

### Quantitative PCR of *CsPLTs*

RNA samples were extracted (Yaffe *et al*., 2012) from hand-dissected 6 dpg ‘9930’ cucumber seedlings grown in square petri dishes on top of a nylon mesh covering a layer of wet potting soil in a long day growth chamber (16 h light, 8 h dark) at 22 °C under white ∼100 μmol/s LED lights. Each of the four biological replicates consisted of pooled tissues of 7 seedlings. Roots were subdivided in four regions, i.e. the bottom ∼ 2 mm containing the apical meristem and the lower elongation zone, and three subsequent ∼3 mm segments encompassing the remaining elongation zone, maturation zone with emerging root hairs, and mature root tissue. A ∼ 3 mm central segment was retrieved from the hypocotyl, and a ∼ 3 x 3 mm piece was taken from one of the cotyledons. The first true leaf was removed from the shoot apices. cDNA was synthesized with the RevertAid Reverse Transcriptase system (Fermentas) using oligo(dT)18 primers. qPCRs were performed with SYBR Green in the Bio-Rad CFX Connect Real-Time PCR Detection System using 3 technical replicates per sample. Primers are listed in Table S6. Cq values were normalised against the *CsUBC* gene (CsaV3_2G031000).

### *In situ* hybridisation

’9930’ cucumbers were grown for 5 days in wet germination pouches (CYG) in a long day growth chamber. Shoot apices were collected in ice-cold fixative (4% paraformaldehyde with 0.5% EM-grade glutaraldehyde in PBS), exposed to repeated 5-minute vacuum bursts until sinking to the bottom of the vial, then incubated at 4°C overnight. Further dehydration, paraffin infiltration and embedding steps are described in (Kulikova *et al*., 2018). The tissues were sectioned longitudinally (20 μm) or transversally (10 μm) prior to *in situ* hybridisation with the Invitrogen™ ViewRNA™ ISH Tissue Assay kit (ThermoFisher Scientific) according to manufacturer’s user guide. Type 1 RNA ISH probe sets were designed from full-length cDNA and synthesized at ThermoFisher Scientific. Catalogue numbers are VPGZFCU for *CsPLT3/7* and VP2W7TZ for *CsPLT5.* Sections were imaged with a Nikon Eclipse 80i using brightfield and fluorescent (TRITC; excitation 527-552 nm, emission 577-632 nm) channels.

### EMS mutagenesis and TILLING

EMS mutagenesis was performed on the ‘long’-type parental inbred line. Seeds were treated with 0.7% EMS and sown on the following day, transplanted after 5 days to pots and transplanted to soil in an unlighted greenhouse 4 weeks after sowing. Plants were insect pollinated and upon maturity fruits were harvested in bulk. For TILLING screens, 5-day old plants were sampled and genotyped by amplicon sequencing (Illumina MiSeq). For selection and seed production in subsequent generations, allele-specific KASP assays were developed.

### Confocal microscopy

6 dpg wild type and *Csplt3/7^Q156*^* shoot apices were collected in Renaissance (RS2200) solution (Musielak *et al*., 2015) and stored at 4 °C until imaging. Tissues were imaged on a Zeiss LSM710 confocal microscope with an excitation wavelength of 405 nm. Prior to imaging with a water dipping lens, SAMs were positioned upright in 3% agarose and submerged in Milli-Q water.

### Greenhouse measurements

Seeds were sown in a greenhouse compartment on rockwool trays, covered with vermiculite, watered with tap water and incubated at 27 °C under natural lighting for 4 days. At this point, germinated plants were sampled for genotyping if required. Sampled plants were stored until genotype selection at 15 °C or directly transplanted to a greenhouse at 23 °C in rockwool blocks with supplemental lighting (Philips Son-T gas pressure lamps, 150 µmol/m^2^s). Plants were transferred to a 22/18 °C (light/dark hours) greenhouse with supplemental lighting (Philips Son-T gas pressure lamps, 150 µmol/m^2^s) 4 weeks past germination, planted in rock wool mats with an automatic watering system and guided upwards along a vertical wire in an umbrella cultivation system. To minimise stem torsion, shoots were attached to the wire using metal rings. Axillary shoots were pruned weekly from the plants, unless counted. Fruits were maintained intermittently (i.e. one fruit per two nodes). Internode length and plant height were measured with a flexible measuring rope. Leaf production per week was determined by weekly attachment of plastic rings around the highest possible internode, then counting the leaves in between successive rings. Axillary shoot counting was performed from top to bottom to account for the initial SAM loss of *Csplt3/7^Q156*^*. Only distinct axillary shoots (> 3 cm) were considered. Phyllotaxis was measured with 360-degree hinged protractors that can be clipped around the cucumber stem, simultaneously allowing measurement of both meristem chirality and angle in 22.5° bins. Chirality was attributed based on the most frequent direction of turning in the upper five measured internodes. Torsion was measured in internode photographs and defined as the angle deflection to the right (counterclockwise) or left (clockwise) from the vertical internode axis. The predicted effect of torsion was calculated according to equation 1 (Landrein et al., 2013):

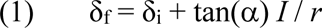

In which δ_f_ is the final divergence angle, δ_i_ the initial divergence angle at the IM, α the torsion angle, *I* internode length, and *r* stem radius.

### RNA-sequencing

RNA-seq was carried out in triplicate on shoot apices dissected from 6 dpg ‘9930’ seedlings grown under long day conditions at 22 °C in soil plates on top of a nylon mesh (7 pooled apices per replicate) and grown in wet germination pouches (20 pooled apices per replicate). The first true leaf from each apex was removed. Tissue was collected in 200 µL RNAlater (Invitrogen) on ice, which was removed after adding 750 µL Milli-Q water prior to freezing in liquid N_2_. RNA was extracted with the LogSpin protocol (Yaffe *et al*., 2012), treated with DNaseI (Qiagen), then purified with EtOH purification. The libraries were sequenced on a NovaSeq6000 Plus at GenomeScan. Data analysis was performed by trimming the reads with fastp v0.23.4 (Chen *et al*., 2018), then pseudo-aligning to the PLAZA5 instance of the full ‘Chinese long 9930 v3’ transcriptome (Van Bel *et al*., 2022) with kallisto v0.46.1 (Bray *et al*., 2016). Genes were considered expressed if TPM > 1 in at least three of the six samples.

### Cloning

The DAP-seq construct *pSPUTK-GG 3xFLAG-GFP* was described previously (Kerstens *et al*., 2024). The similar constructs *pSPUTK-GG 3xFLAG-CsPLT3/7* and *pSPUTK-GG 3xFLAG-CsPLT5* were assembled by combining *pSPUTK-GG* (Kerstens *et al*., 2024), *pICSL30005* (Addgene #50299) and a BsaI-cCsPLT3/7 or BsaI-cCsPLT5 amplicon through BsaI cloning. Transcriptional reporters were generated by amplifying 6.1 kb (*CsPLT1/2*) or 6.3 kb (*CsPLT3/7*) promoter + 5’UTR fragments from ‘9930’ gDNA and subsequent Gateway cloning into pGEM-T Easy 221 (Invitrogen) and pGWB3 (Nakagawa *et al*., 2007). Arabidopsis was transformed by floral dip (Clough & Bent, 1998).

### DAP-sequencing

To obtain genomic DNA with SAM-specific modifications, dissected SAMs without the first true leaves were collected from 6 dpg parental inbred line seedlings grown in wet germination pouches (CYG) in a long day (18h light, 6h dark) greenhouse with fluctuating temperatures (day: 20.5 °C; night: 16.0 °C). Two replicates of 100 SAMs each were collected on ice and snap-frozen in liquid N_2_. Further DAP and analysis steps were carried out as described previously (Kerstens *et al*., 2025a), using the ‘Chinese long 9930 v3’ genome (Li *et al*., 2019) and integrating replicates through IDR analysis (https://github.com/nboley/idr). Fragments of reads in peaks (FRiP) was calculated by dividing the number of filtered, mapped reads within peaks by the total number of filtered, mapped reads per sample. (Co-)orthology of target genes was inferred from both collinearity (’anchor point’) and best BLAST hits (’BHIF’) as available from the Integrative Orthology data within the PLAZA5 Dicots database (Van Bel *et al*., 2022).

### GUS staining

Arabidopsis Col-0 plants were grown for 5-6 weeks in long day conditions (16 h light, 8 h dark) at 22 °C under white LED lights. Ovules were hand-dissected from young siliques and directly transferred to GUS staining solution (50 mM sodium phosphate buffer pH 7.0, 2 mM potassium ferro/ferricyanide, 0.1% IGEPAL CA-630, 0.5 mg/mL X-GlcA). Submerged ovules were vacuum infiltrated for 15 minutes, then stained overnight at 37 °C. Stained ovules were cleared at 4 °C in chloral hydrate solution (8 g chloral hydrate, 1 g glycerol, 3 mL H_2_O) and imaged using a Zeiss Axio Imager A1.

## Author contributions

Conceptualization: M.K., B.S., V.W.; Methodology: M.K., B.S., V.W.; Investigation: M.K., F.M., K.A., O.K., M.L.; Writing: M.K., V.W.; Editing: B.S.; Visualization: M.K.; Supervision: B.S., V.W.; Project administration: V.W.; Funding acquisition: M.K.

## Significance statement

We reveal that shoot apical meristem-expressed *PLETHORA3/7* in cucumber shapes shoot architecture. *Csplt3/7* mutant mature embryos initially lack shoot apical meristems, but later form secondary meristems from their apex, resulting in shoots with irregular internode length, phyllotaxis, and aberrant development of organs in leaf axils.

## Acknowledgements

We thank Freek van der Klugt, Rory McCarthy and Jarno van Schijndel for assisting with greenhouse measurements, wet lab experiments, and amplification and size-selection of DAP libraries. Taco Jesse is acknowledged for coordination of RNA and DAP-sequencing, and Kees van Dun for stimulating discussions.

## Data availability

The DAP-seq and RNA-seq data are available in the Gene Expression Omnibus with accession numbers GSE294326 and GSE294327, respectively.

## References

Aida M, Beis D, Heidstra R, Willemsen V, Blilou I, Galinha C, Nussaume L, Noh YS, Amasino R, Scheres B. 2004. The PLETHORA genes mediate patterning of the Arabidopsis root stem cell niche. Cell 119: 109–120.

Aida M, Ishida T, Fukaki H, Fujisawa H, Tasaka M. 1997. Genes involved in organ separation in Arabidopsis: an analysis of the cup-shaped cotyledon mutant. The Plant Cell 9: 841–857.

Bahafid E, Bradtmöller I, Thies AM, Ton Nguyen T, Gutierrez C, Desvoyes B, Stahl Y, Blilou I, Simon RGW. 2023. The Arabidopsis SHORTROOT network coordinates shoot apical meristem development with auxin-dependent lateral organ initiation. eLife 12.

Barrera-Redondo J, Ibarra-Laclette E, Vázquez-Lobo A, Gutiérrez-Guerrero YT, Sánchez de la Vega G, Piñero D, Montes-Hernández S, Lira-Saade R, Eguiarte LE. 2019. The Genome of Cucurbita argyrosperma (Silver-Seed Gourd) Reveals Faster Rates of Protein-Coding Gene and Long Noncoding RNA Turnover and Neofunctionalization within Cucurbita. Molecular Plant 12: 506–520.

Barton MK, Poethig RS. 1993. Formation of the shoot apical meristem in Arabidopsis thaliana: an analysis of development in the wild type and in the shoot meristemless mutant. Development 119: 823–831.

Belles-Boix E, Hamant O, Witiak SM, Morin H, Traas J, Pautot V. 2006. KNAT6: An Arabidopsis Homeobox Gene Involved in Meristem Activity and Organ Separation. The Plant Cell 18: 1900–1907.

Besnard F, Refahi Y, Morin V, Marteaux B, Brunoud G, Chambrier P, Rozier F, Mirabet V, Legrand J, Lainé S, et al. 2014. Cytokinin signalling inhibitory fields provide robustness to phyllotaxis. Nature 505: 417–421.

Brand U, Fletcher JC, Hobe M, Meyerowitz EM, Simon R. 2000. Dependence of Stem Cell Fate in Arabidopsis on a Feedback Loop Regulated by CLV3 Activity. Science 289: 617–619.

Bray NL, Pimentel H, Melsted P, Pachter L. 2016. Near-optimal probabilistic RNA-seq quantification. Nature Biotechnology 34: 525–527.

Capella-Gutiérrez S, Silla-Martínez JM, Gabaldón T. 2009. trimAl: A tool for automated alignment trimming in large-scale phylogenetic analyses. Bioinformatics 25: 1972–1973.

Che G, Gu R, Zhao J, Liu X, Song X, Zi H, Cheng Z, Shen J, Wang Z, Liu R, et al. 2020. Gene regulatory network controlling carpel number variation in cucumber. Development 147: dev184788.

Chen B, Maas L, Figueiredo D, Zhong Y, Reis R, Li M, Horstman A, Riksen T, Weemen M, Liu H, et al. 2022. BABY BOOM regulates early embryo and endosperm development. Proceedings of the National Academy of Sciences of the United States of America 119.

Chen S, Zhou Y, Chen Y, Gu J. 2018. Fastp: An ultra-fast all-in-one FASTQ preprocessor. Bioinformatics 34: i884–i890.

Cheng F, Song M, Zhang M, Cheng C, Chen J, Lou Q. 2022. A SNP mutation in the CsCLAVATA1 leads to pleiotropic variation in plant architecture and fruit morphogenesis in cucumber (Cucumis sativus L.). Plant Science 323: 111397.

Clark SE, Running MP, Meyerowitz EM. 1993. CLAVATA1, a regulator of meristem and flower development in Arabidopsis. Development 119: 397–418.

Clark SE, Running MP, Meyerowitz EM. 1995. CLAVATA3 is a specific regulator of shoot and floral meristem development affecting the same processes as CLAVATA1. Development 121: 2057–2067.

Clough SJ, Bent AF. 1998. Floral dip: A simplified method for Agrobacterium-mediated transformation of Arabidopsis thaliana. Plant Journal 16: 735–743.

DeYoung BJ, Bickle KL, Schrage KJ, Muskett P, Patel K, Clark SE. 2006. The CLAVATA1-related BAM1, BAM2 and BAM3 receptor kinase-like proteins are required for meristem function in Arabidopsis. The Plant Journal 45: 1–16.

Du Y, Scheres B. 2017. PLETHORA transcription factors orchestrate de novo organ patterning during Arabidopsis lateral root outgrowth. Proceedings of the National Academy of Sciences of the United States of America 114: 11709–11714.

Galinha C, Hofhuis H, Luijten M, Willemsen V, Blilou I, Heidstra R, Scheres B. 2007. PLETHORA proteins as dose-dependent master regulators of Arabidopsis root development. Nature 449: 1053–1057.

Galvan-Ampudia CS, Cerutti G, Legrand J, Brunoud G, Martin-Arevalillo R, Azais R, Bayle V, Moussu S, Wenzl C, Jaillais Y, et al. 2020. Temporal integration of auxin information for the regulation of patterning (J Kleine-Vehn and CS Hardtke, Eds.). eLife 9: e55832.

Han L, Huang Y, Li C, Tian D, She D, Li M, Wang Z, Chen J, Liu L, Wang S, et al. 2024. Heterotrimeric Gα-subunit regulates flower and fruit development in CLAVATA signaling pathway in cucumber. Horticulture Research 11: uhae110.

Heisler MG, Ohno C, Das P, Sieber P, Reddy G V, Long JA, Meyerowitz EM. 2005. Patterns of Auxin Transport and Gene Expression during Primordium Development Revealed by Live Imaging of the Arabidopsis Inflorescence Meristem. Current Biology 15: 1899–1911.

Jönsson H, Heisler MG, Shapiro BE, Meyerowitz EM, Mjolsness E. 2006. An auxin-driven polarized transport model for phyllotaxis. Proceedings of the National Academy of Sciences 103: 1633–1638.

Kareem A, Durgaprasad K, Sugimoto K, Du Y, Pulianmackal AJ, Trivedi ZB, Abhayadev P V., Pinon V, Meyerowitz EM, Scheres B, et al. 2015. PLETHORA genes control regeneration by a two-step mechanism. Current Biology 25: 1017–1030.

Katoh K, Rozewicki J, Yamada KD. 2017. MAFFT online service: multiple sequence alignment, interactive sequence choice and visualization. Briefings in Bioinformatics bbx108.

Kerstens M, Galinha C, Hofhuis H, Nodine M, Pardal R, Scheres B, Willemsen V. 2024. PLETHORA transcription factors promote early embryo development through induction of meristematic potential. Development 151: dev202527.

Kerstens MHL, van der Klugt F, Hofhuis H, Scheres B, Willemsen V. 2025a. PLETHORAs shape Arabidopsis phyllotaxis through modulation of patterning robustness and accelerated inflorescence development. New Phytologist Manuscript accepted.

Kerstens MHL, Schranz ME, Bouwmeester K. 2020. Phylogenomic analysis of the APETALA2 transcription factor subfamily across angiosperms reveals both deep conservation and lineage-specific patterns. Plant Journal 103: 1516–1524.

Kulikova O, Franken C, Bisseling T. 2018. In Situ Hybridization Method for Localization of mRNA Molecules in Medicago Tissue Sections. In: Cañas LA, Beltrán JP, eds. Functional Genomics in Medicago truncatula: Methods and Protocols. New York, NY: Springer New York, 145–159.

Landrein B, Lathe R, Bringmann M, Vouillot C, Ivakov A, Boudaoud A, Persson S, Hamant O. 2013. Impaired Cellulose Synthase Guidance Leads to Stem Torsion and Twists Phyllotactic Patterns in Arabidopsis. Current Biology 23: 895–900.

Landrein B, Refahi Y, Besnard F, Hervieux N, Mirabet V, Boudaoud A, Vernoux T, Hamant O. 2015. Meristem size contributes to the robustness of phyllotaxis in Arabidopsis. Journal of Experimental Botany 66: 1317–1324.

Laux T, Mayer KFX, Berger J, Jürgens G. 1996. The WUSCHEL gene is required for shoot and floral meristem integrity in Arabidopsis. Development 122: 87–96.

Letunic I, Bork P. 2019. Interactive Tree Of Life (iTOL) v4: recent updates and new developments. Nucleic Acids Research 47: W256–W259.

Li Q, Li H, Huang W, Xu Y, Zhou Q, Wang S, Ruan J, Huang S, Zhang Z. 2019. A chromosome-scale genome assembly of cucumber (Cucumis sativus L.). GigaScience 8: giz072.

Liu X, Chen J, Zhang X. 2021. Genetic regulation of shoot architecture in cucumber. Horticulture Research 8: 143.

Liu Q, Han D, Chen J, Wang J, Cheng D, Chen X, Jiang J, Tian S, Wang J, Liu M, et al. 2025. ClBBM and ClPLT2 function redundantly during both male and female gametophytes development in watermelon. Horticultural Plant Journal 11: 323–335.

Ma L, Wang Q, Zheng Y, Guo J, Yuan S, Fu A, Bai C, Zhao X, Zheng S, Wen C, et al. 2022. Cucurbitaceae genome evolution, gene function, and molecular breeding. Horticulture Research 9: uhab057.

Mayer KFX, Schoof H, Haecker A, Lenhard M, Jürgens G, Laux T. 1998. Role of WUSCHEL in Regulating Stem Cell Fate in the Arabidopsis Shoot Meristem. Cell 95: 805– 815.

Montero-Pau J, Blanca J, Bombarely A, Ziarsolo P, Esteras C, Martí-Gómez C, Ferriol M, Gómez P, Jamilena M, Mueller L, et al. 2018. De novo assembly of the zucchini genome reveals a whole-genome duplication associated with the origin of the Cucurbita genus. Plant Biotechnology Journal 16: 1161–1171.

Mudunkothge JS, Krizek BA. 2012. Three Arabidopsis AIL/PLT genes act in combination to regulate shoot apical meristem function. Plant Journal 71: 108–121.

Müller R, Borghi L, Kwiatkowska D, Laufs P, Simon R. 2006. Dynamic and Compensatory Responses of Arabidopsis Shoot and Floral Meristems to CLV3 Signaling. The Plant Cell 18: 1188–1198.

Musielak TJ, Schenkel L, Kolb M, Henschen A, Bayer M. 2015. A simple and versatile cell wall staining protocol to study plant reproduction. Plant Reproduction 28: 161–169.

Nakagawa T, Kurose T, Hino T, Tanaka K, Kawamukai M, Niwa Y, Toyooka K, Matsuoka K, Jinbo T, Kimura T. 2007. Development of series of gateway binary vectors, pGWBs, for realizing efficient construction of fusion genes for plant transformation. Journal of Bioscience and Bioengineering 104: 34–41.

Nguyen LT, Schmidt HA, Von Haeseler A, Minh BQ. 2015. IQ-TREE: A fast and effective stochastic algorithm for estimating maximum-likelihood phylogenies. Molecular Biology and Evolution 32: 268–274.

Pinon V, Prasad K, Grigg SP, Sanchez-Perez GF, Scheres B. 2013. Local auxin biosynthesis regulation by PLETHORA transcription factors controls phyllotaxis in Arabidopsis. Proceedings of the National Academy of Sciences of the United States of America 110: 1107– 1112.

Prasad K, Grigg SP, Barkoulas M, Yadav RK, Sanchez-Perez GF, Pinon V, Blilou I, Hofhuis H, Dhonukshe P, Galinha C, et al. 2011. Arabidopsis PLETHORA transcription factors control phyllotaxis. Current Biology 21: 1123–1128.

Reddy GV, Heisler MG, Ehrhardt DW, Meyerowitz EM. 2004. Real-time lineage analysis reveals oriented cell divisions associated with morphogenesis at the shoot apex of Arabidopsis thaliana. Development 131: 4225–4237.

Reinhardt D, Pesce E-R, Stieger P, Mandel T, Baltensperger K, Bennett M, Traas J, Friml J, Kuhlemeier C. 2003. Regulation of phyllotaxis by polar auxin transport. Nature.

de Reuille PB, Bohn-Courseau I, Ljung K, Morin H, Carraro N, Godin C, Traas J. 2006. Computer simulations reveal properties of the cell-cell signaling network at the shoot apex in Arabidopsis. Proceedings of the National Academy of Sciences 103: 1627–1632.

Robertson C, Xue H, Saltini M, Fairnie ALM, Lang D, Kerstens MHL, Willemsen V, Ingle RA, Barrett SCH, Deinum EE, et al. 2025. Spiral phyllotaxis predicts left-right asymmetric growth and style deflection in mirror-image flowers of Cyanella alba. Nature Communications 16: 3695.

Santuari L, Sanchez-Perez GF, Luijten M, Rutjens B, Terpstra I, Berke L, Gorte M, Prasad K, Bao D, Timmermans-Hereijgers JLPM, et al. 2016. The PLETHORA gene regulatory network guides growth and cell differentiation in Arabidopsis roots. Plant Cell 28: 2937–2951.

Schoof H, Lenhard M, Haecker A, Mayer KFX, Jürgens G, Laux T. 2000. The Stem Cell Population of Arabidopsis Shoot Meristems Is Maintained by a Regulatory Loop between the CLAVATA and WUSCHEL Genes. Cell 100: 635–644.

Smith RS, Guyomarc’h S, Mandel T, Reinhardt D, Kuhlemeier C, Prusinkiewicz P. 2006. A plausible model of phyllotaxis. Proceedings of the National Academy of Sciences.

Stuurman J, Jäggi F, Kuhlemeier C. 2002. Shoot meristem maintenance is controlled by a GRAS-gene mediated signal from differentiating cells. Genes & Development 16: 2213–2218.

Sun H, Wu S, Zhang G, Jiao C, Guo S, Ren Y, Zhang J, Zhang H, Gong G, Jia Z, et al. 2017. Karyotype Stability and Unbiased Fractionation in the Paleo-Allotetraploid Cucurbita Genomes. Molecular Plant 10: 1293–1306.

Van Bel M, Silvestri F, Weitz EM, Kreft L, Botzki A, Coppens F, Vandepoele K. 2022. PLAZA 5.0: extending the scope and power of comparative and functional genomics in plants. Nucleic Acids Research 50: D1468–D1474.

Wang B, Smith SM, Li J. 2018. Genetic Regulation of Shoot Architecture. Annual Review of Plant Biology 69: 437–468.

Weigel D, Alvarez J, Smyth DR, Yanofsky MF, Meyerowitz EM. 1992. LEAFY controls floral meristem identity in Arabidopsis. Cell 69: 843–859.

Yaffe H, Buxdorf K, Shapira I, Ein-Gedi S, Moyal-Ben Zvi M, Fridman E, Moshelion M, Levy M. 2012. LogSpin: a simple, economical and fast method for RNA isolation from infected or healthy plants and other eukaryotic tissues. BMC Research Notes 5: 45.

Yu J, Wu S, Sun H, Wang X, Tang X, Guo S, Zhang Z, Huang S, Xu Y, Weng Y, et al. 2023. CuGenDBv2: an updated database for cucurbit genomics. Nucleic Acids Research 51: D1457–D1464.

Zebell SG, Martí-Gómez C, Fitzgerald B, Cunha CP, Lach M, Seman BM, Hendelman A, Sretenovic S, Qi Y, Bartlett M, et al. 2025. Cryptic variation fuels plant phenotypic change through hierarchical epistasis. Nature 644: 984–992.

Zhao W, Chen Z, Liu X, Che G, Gu R, Zhao J, Wang Z, Hou Y, Zhang X. 2018. CsLFY is required for shoot meristem maintenance via interaction with WUSCHEL in cucumber (Cucumis sativus). New Phytologist 218: 344–356.

