## Supplemental figures for "A PLETHORA transcription factor shapes cucumber shoot architecture"

### Supporting information

#### Supporting table legends

**Table S1.** CsPLT3/7 and CsPLT5 DAP-seq peaks and their position relative to the nearest TSS.

**Table S2**. DAP-seq experimental metrics.

**Table S3.** Genes expressed in the cucumber SAM in 6 dpg seedlings grown on soil plates or in germination pouches (TPM > 1 in at least 3/6 samples). Values are TPM.

**Table S4.** Cucumber SAM-expressed genes with a DAP peak in a [-5 kbp, 5 kbp] range from the TSS and their Arabidopsis orthologs**.**

**Table S5**. TILLING *Csplt* alleles generated in this study.

**Table S6**. Oligonucleotides used in this study.

**Supporting figures**

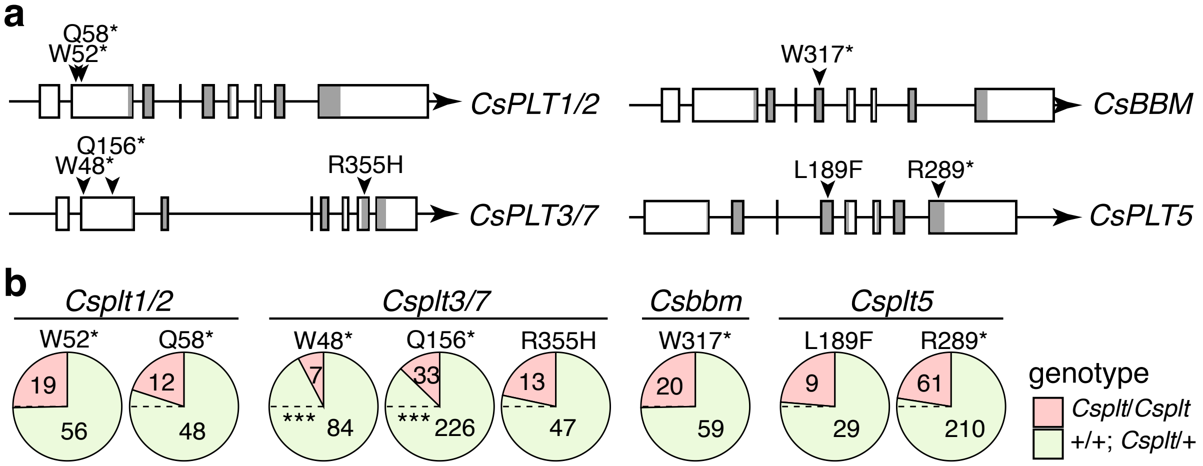


**Figure S1**. *Csplt3/7* premature stop mutants are underrepresented in the progeny of a selfed heterozygous line. (**a**) Gene models displaying the location and nature of the TILLING mutations. AP2 domains indicated in grey. (**b**) Pie charts displaying the proportion of homozygous mutants (red) and heterozygous + wild type seedlings (green) per analysed TILLING allele, with counts indicated. The dashed line indicates the expected frequency of 0.25 based on Mendelian segregation. Statistics are Benjamini-Hochberg and Yates-corrected z-tests (freq. < 0.25; *** left: p = 4.5e-4; right: p = 2.9e-5).

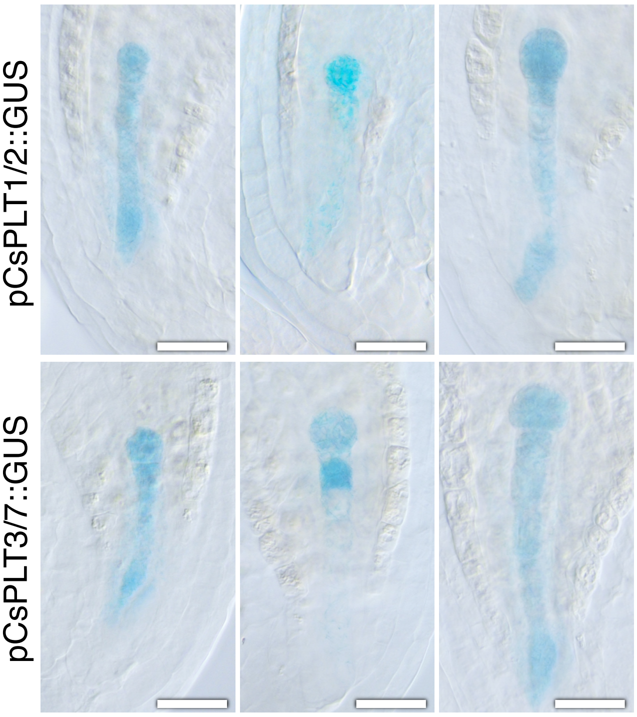


**Figure S2**. *pCsPLT1/2* and *pCsPLT3/7* are active in the Arabidopsis embryo. Shown are DIC images of overnight GUS staining of 1-cell stage embryos apical and basal cell expression (left), 1/2-cell stage embryos with predominant apical expression (middle), and 2/4-cell stage embryos with embryo proper and suspensor expression (right). Scale bars are 25 µm.

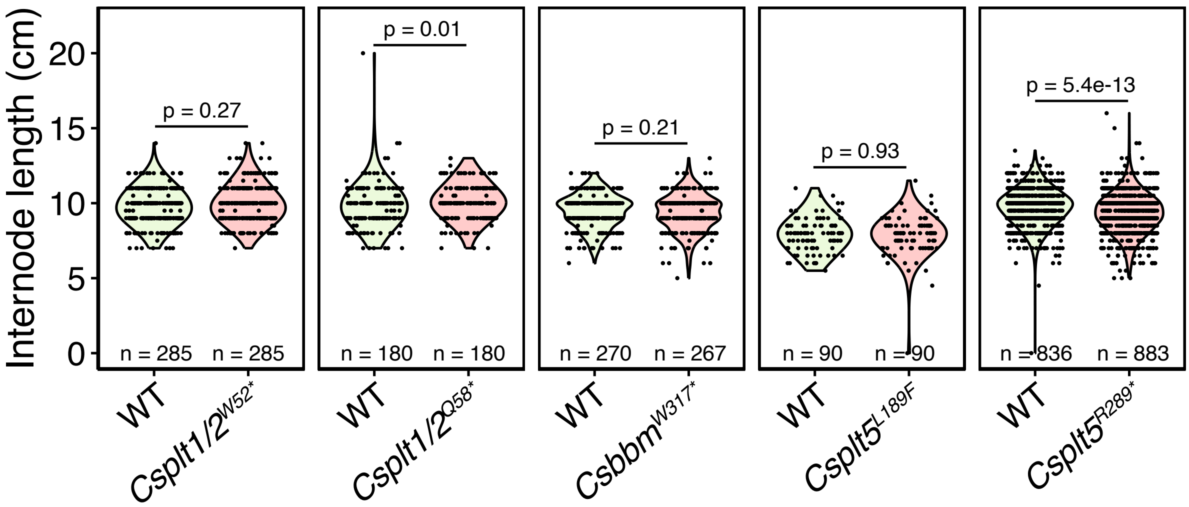


**Figure S3.** *Csplt1/2*, *Csbbm* and *Csplt5* mutants have no shoot phenotype. Internode length of wild type (WT) and mutant progeny from the same selfing, measured with 0.5 cm accuracy. Significance from two-tailed Wilcoxon rank-sum tests.

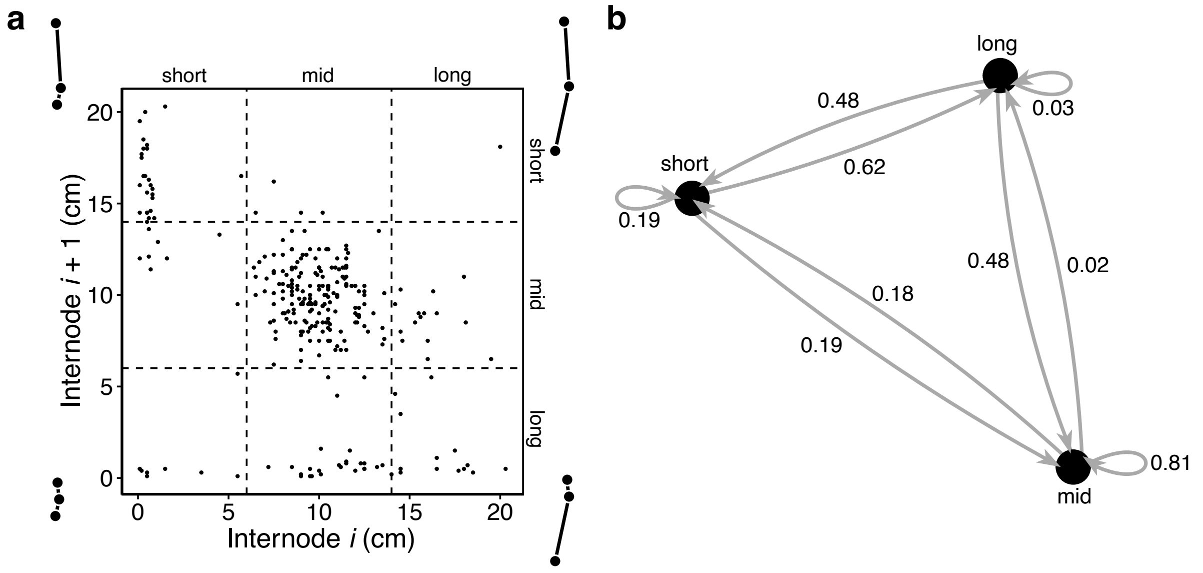


**Figure S4.** *Csplt3/7*$^{Q156*}$ long and short internodes preferentially succeed each other. (**a**) Scatterplot showing the relationship between the length of internode *i* (x-axis) and its succeeding internode *i + 1* (y-axis). Dotted lines indicate the classes defined as short (≤ 6 cm), mid (> 6 & < 14 cm), and long (≥ 14 cm). Ball-and-stick diagrams schematically represent the node (ball) and internode (stick) conformation in the corners of the plot. (**b**) Markov chain diagram showing probabilities of transitioning between the three internode classes.

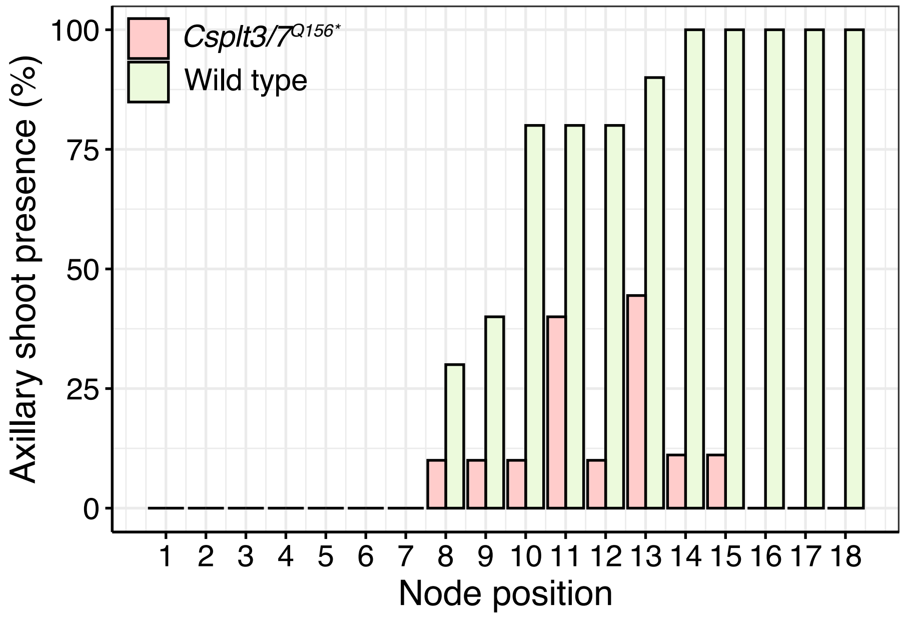


**Figure S5.** *Csplt3/7*$^{Q156*}$ axillary shoots fail to form consistently. Bars indicate percentage of nodes, counted from the apex of 10 wild type and mutant plants, that formed axillary shoots in 2-month-old plants. Note that one mutant plant only had 12 nodes at this age, thus nodes 13 to 18 are based on 9 mutant plants.

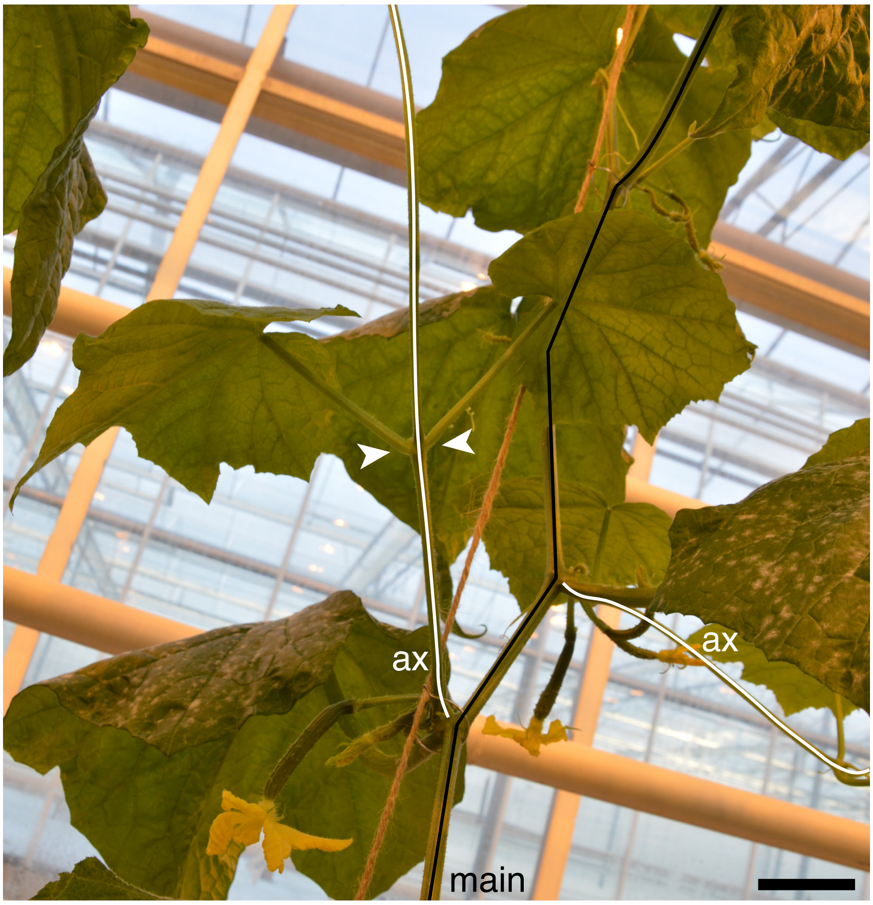


**Figure S6.** *Csplt3/7*$^{Q156*}$ axillary shoot (ax; white) internode length mimics the main shoot (main; black) phenotype. Arrowheads point to near-co-initiated leaves, which is succeeded by a very long internode. Scale bar is 5 cm.

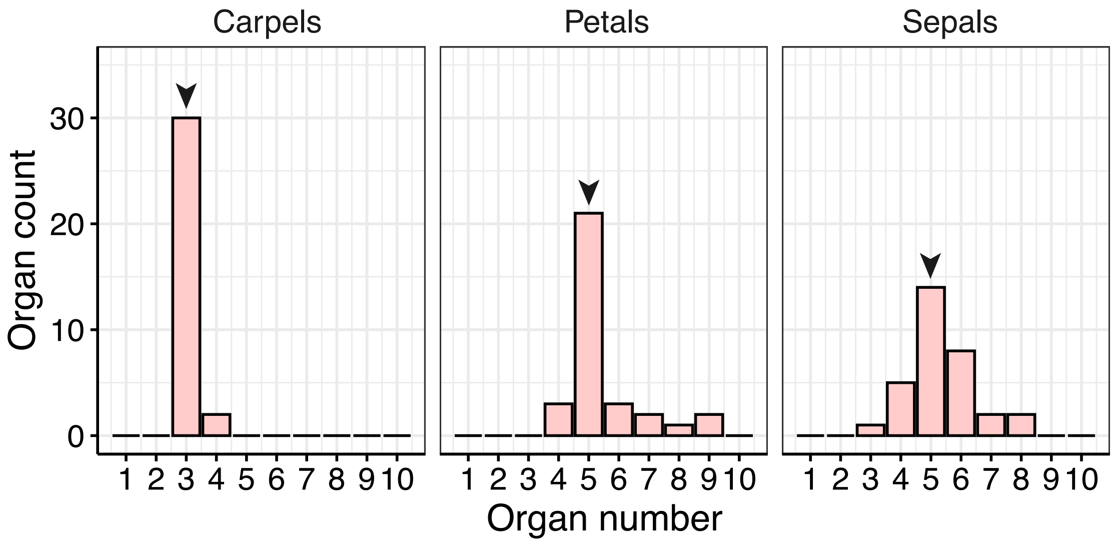


**Figure S7.** Floral organ counts of *Csplt3/7*$^{Q156*}$ (n = 32). Arrows indicate the invariant counts of wild type flowers.

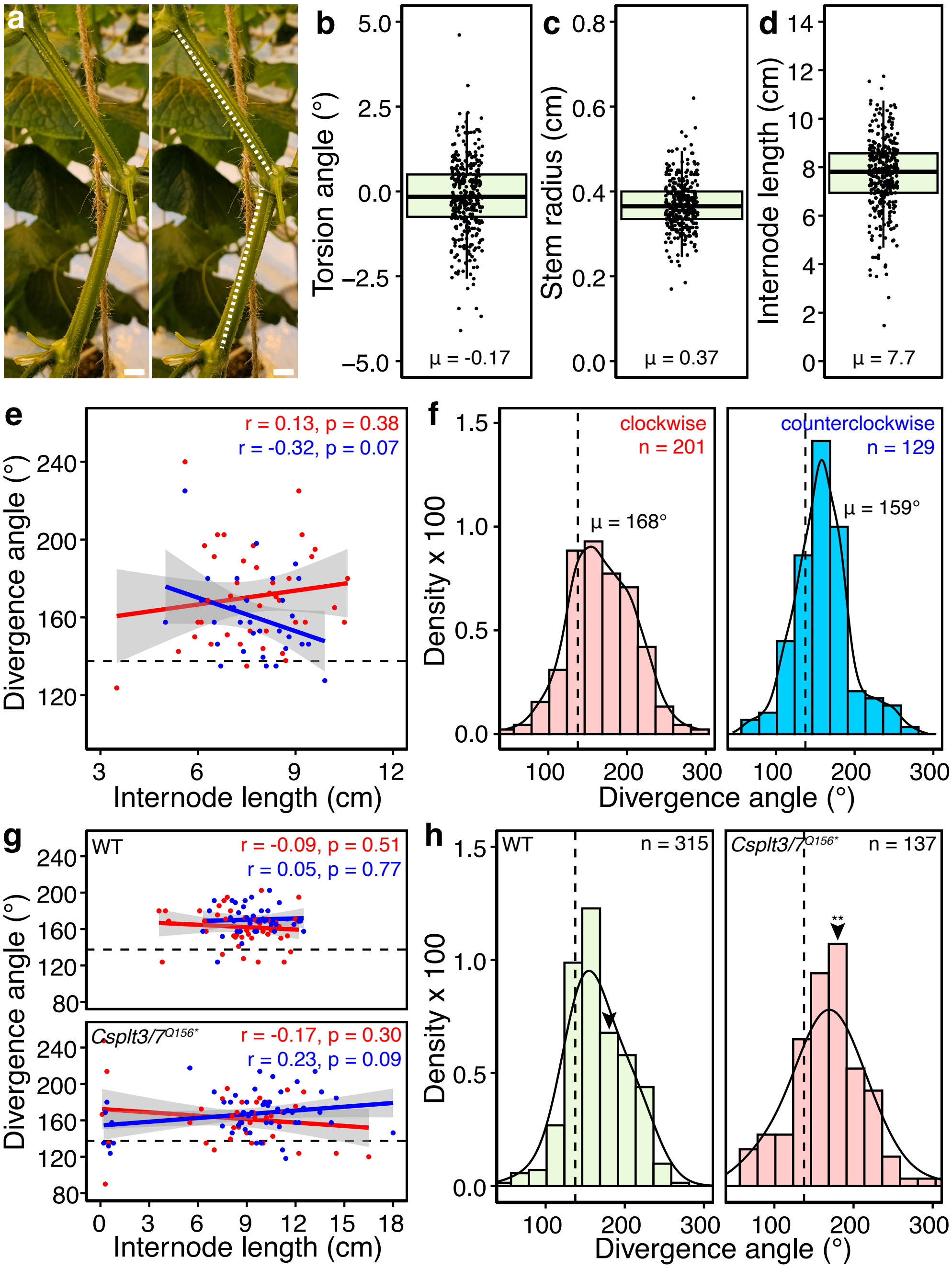


**Figure S8**. Cucumber phyllotaxis is not consistently chirality-dependently modified by stem torsion. (**a**) Two internodes exhibiting clockwise stem torsion. Dashed lines (right) are positioned on top of a stem ridge. Scale bar is 1 cm. (**b-d**) Measured values for torsion angle, stem radius and internode length of 350 internodes originating from 15 WT plants. (**e**) Leaf divergence angles against internode length in clockwise (red) and counterclockwise (blue) meristems of wild type plants. Each dot corresponds to the average value of at least two measurements in 0.1 cm bins. Correlation coefficient and significance of the linear regressions are indicated. Shaded areas correspond to the 95% confidence interval. (**f**) Divergence angle histograms and density plot of clockwise (left; red) and counterclockwise (right; blue) meristems in 22.5° bins. (**g**) Leaf divergence angles against internode length of *Csplt3/7*$^{Q156*}$ and its wild type sister progeny (at least 2 values per 0.1 cm bin). (**h**) Histograms and density plots displaying the occurrence of divergence angles of same-size internodes (9.0 to 11.0 cm). Arrowheads points towards the bin containing 180°. Significance from a Y-corrected z-test ($p = 3.7e-3$). The dashed line in (**e**-**h**) corresponds to the golden angle of 137.5°.
